# Molecular Determinants of Liquid Demixing and Amyloidogenesis in Human CPEB3

**DOI:** 10.1101/2020.06.02.129783

**Authors:** Daniel Ramírez de Mingo, Paula López-García, Rubén Hervás, Douglas V. Laurents, Mariano Carrión-Vázquez

**Affiliations:** Instituto Cajal, CSIC. Avenida Doctor Arce 37, Madrid 28002, Spain; Stowers Institute for Medical Research, Kansas City, MO 64110, USA; Instituto de Química Física “Rocasolano”, CSIC. C/ Serrano 119, Madrid 28006, Spain

**Keywords:** hCPEB3, memory consolidation, intrinsically disordered protein, liquid-liquid phase separation, functional amyloid

## Abstract

The cytoplasmic polyadenylation element-binding protein 3 (CPEB3), is an RNA-binding protein which in its soluble state is localized in membraneless neuronal RNA granules keeping target mRNAs in a repressed state. The stimulus-dependent aggregation of CPEB3 activates target mRNAs translation, a central event for the maintenance of long-term memory-related synaptic plasticity in mammals. To date, the molecular determinants that govern both connected events remain unclear. Here, to gain insight into these processes, the biophysical properties of the human CPEB3 (hCPEB3) are characterized. We found that hCPEB3 homotypic condensation is mainly driven by hydrophobic interactions and occurs under physiological conditions. Moreover, hCPEB3 biomolecular condensates are dynamic inside living cells, whose localization and stabilization are mediated by its RNA-recognition domains. In contrast, the hCPEB3 polar N-terminal region is crucial for hCPEB3 amyloid-like aggregation *in vitro*, which is disrupted by the polyglutamine binding peptide 1 (QBP1), A*β*_42_ seeds and Hsp70, highlighting the importance of the Q_4_RQ_4_ tract as well as the hydrophobic residues for hCPEB3 functional aggregation. Based on these findings, we postulate a model for hCPEB3’s role in memory persistence that advances a rather sophisticated control for hCPEB3 condensate dissociation and amyloid-like formation to achieve its physiological function.

**Highlights:**

- hCPEB3 forms toxic intermediates that persist longer than in other functional amyloids.
- RNA-recognition domains stabilize hCPEB3 granule formation and dynamics.
- Different segments within hCPEB3 promote amyloidogenesis and liquid demixing.
- hCPEB3 amyloid formation requires both hydrophobic and polyQ segments.

**Graphical Abstract:** 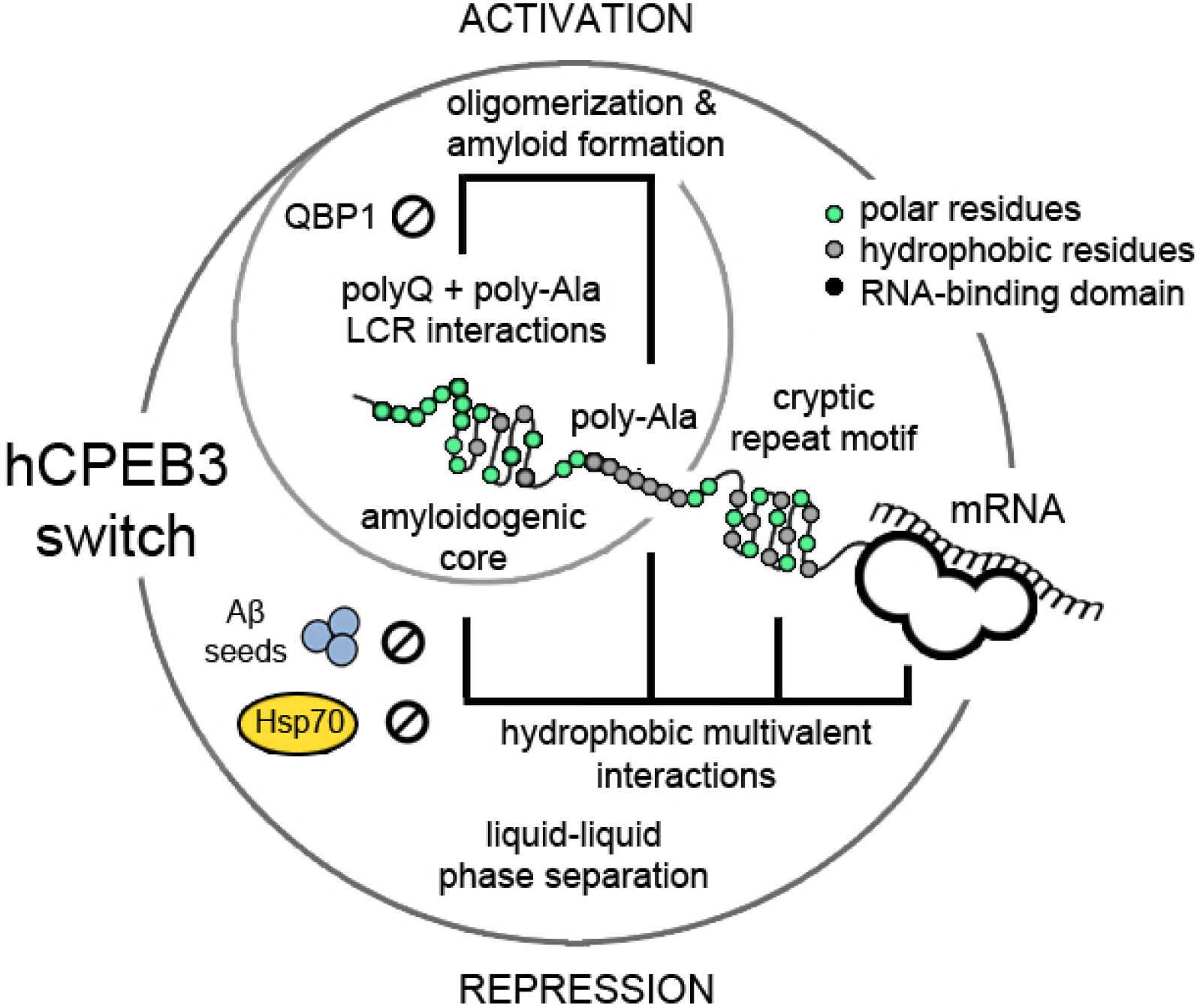

## Introduction

Memory is all you are as a person, an embroidery of your own personal experience you weave throughout your life. The human brain has the ability to encode, retain and subsequently recall previous experiences, and the sum of what we remember we call it memory. Synaptic plasticity, the biological process by which neurons strengthen their connections in discrete brain networks, is key for memory storage and depends on protein synthesis^1^. Recent evidences indicate the impact of the translational control of specific mRNAs on synaptic stabilization during <u>l</u>ong-<u>t</u>erm <u>p</u>otentiation (LTP). However, the underlying molecular mechanisms by which protein translational regulators mediate LTP within synapses are not fully understood. Of particular interest is the <u>c</u>ytoplasmic <u>p</u>olyadenylation <u>e</u>lement-<u>b</u>inding (CPEB) family of prion-like RNA-binding proteins, which comprise four homologs in mammals (CPEB1-4)^2,3^. In the hippocampus, soluble CPEB3 represses the translation of plasticity-related proteins including the scaffolding protein PSD-95, the NMDA receptors, the AMPA receptor subunits GluR1 and GluR2 and β-actin^4–6^. Upon neuronal stimulation, CPEB3 polymerizes into an active, aggregated form that triggers the polyadenylation and derepression of target mRNAs leading to LTP activation^7–9^. Mouse CPEB3 (mCPEB3) forms amyloid filaments *in vitro* through the N-terminal <u>p</u>rion-<u>l</u>ike <u>d</u>omain (PLD), whose properties resemble those of *Aplysia* (*Ap*CPEB) and *Drosophila* (Orb2) homologs^10–13^. Both the mCPEB3 and the human (hCPEB3) homologs, which share 89% of sequence identity, have a PLD that is longer and poorer in Q residues than both Orb2, and particularly *Ap*CPEB. In principle, this different sequence organization may suggest that mCPEB3 and hCPEB3’s prion-like regions may be more tightly regulated to afford a finer control of memory consolidation in mammals. Moreover, the conformational trends within the PLD that precedes amyloid formation appears to be different among species. Thus, while the *Ap*CPEB PLD forms *α*-helical coiled-coils *in vitro* and assembles into multimers to enhance amyloid aggregation^14–16^, the recombinant Orb2 PLD remains as a random coil conformation, which is able to oligomerize and form amyloid structures whose *β*-sheet core seems not to include the Q-rich domain^17^. By contrast, recent cryo-EM data revealed that *Drosophila* Orb2 from adult heads forms a threefold-symmetric hydrophilic amyloid core based on the Q-rich stretch^18^. In spite of these differences, experimental data suggest that *Ap*CPEB, Orb2 and mCPEB3 adopt an amyloid-like state that plays a key role in memory consolidation^6,12,15^.

Eukaryotic cells contain numerous membraneless organelles, also called biomolecular condensates. These organelles compartmentalize a myriad of specific proteins and/or nucleic acids *via* <u>l</u>iquid-<u>l</u>iquid <u>p</u>hase <u>s</u>eparation (LLPS), which enables a strict spatial and temporal control of the cellular biochemical reactions^19,20^. In the last years, the role of post-translational modifications and the nuclear transport system on phase transition regulation has started to be elucidated for intrinsically disordered proteins (IDPs) with PLDs such as FUS or TDP-43, which are RNA-binding proteins with similar organization than hCPEB3^21,22^. Current evidence supports that, in contrast to Orb2^23^, mCPEB3 is SUMOylated in the basal state and represses translation of target mRNAs by retaining them into neuronal RNA granules through LLPS^24^. Furthermore, mCPEB3 is phosphorylated by protein kinase A and calcium/calmodulin-dependent protein kinase II, which shuttles in and out of the nucleus^25,26^. However, the importance of post-translational modification sites as well as the leucine-rich nuclear export signal (NES, L349-L353) in hCPEB3 phase separation and amyloid formation is not yet clear. Indeed, the molecular determinants within the CPEB3 sequence that contribute to its physiological function are unknown. Thus, the elucidation of the biophysical eatures that govern hCPEB3 demixing and amyloidogenesis and their association with synaptic plasticity is essential for a full understanding of the molecular bases of memory.

Here, by using a “divide and conquer” strategy based on a serial deletion analysis, we report that the intrinsically disordered region (IDR) of hCPEB3 spontaneously condensates *via* LLPS in physiological conditions and forms a solid amyloid-like state by gelation. We found that hCPEB3’s PLD possesses three key elements for aggregation: the N-terminal Q_4_RQ_4_ polyQ tract, essential for oligomerization; a hydrophobic core, within residues 106-165; and an aggregation-promoting poly-Ala stretch, which interact and collaborate to form the hCPEB3 amyloid. In addition, the non-compositionally bias hCPEB3 IDR’s C-terminal subsequences, but not the PLD, are sufficient and necessary for hCPEB3 IDR LLPS and modulate the kinetics of amyloid assembly *in vitro*. Finally, a mechanistic insight into how hCPEB3 LLPS is driven by hydrophobic intermolecular interactions that can be blocked by both the chaperone Hsp70 and A*β*_42_ is also provided. Our results shed light into how hCPEB3 assembles into macromolecular structures, such as the neuronal granules, and contribute to the understanding of the molecular forces governing hCPEB3 LLPS and amyloidogenesis, which are key events in memory stabilization in mammals. In addition to allow us to understand how protein assembly is orchestrated in the cellular context to avoid toxicity, this knowledge may help us to identify new pharmacological targets for developing new therapeutic strategies for several fatal, amyloid-based diseases^27^.

## Results

### hCPEB3 IDR organization: structural elements with different composition and complexity

An *in silico* analysis of the hCPEB3 sequence, using IUPred^28^, predicted that the N-terminal IDR, spanning residues 1-426, possesses a low net charge with high tendency to adopt a disorder state, in agreement with our recent NMR data^29^ (Fig. 1A). The subsequences with propensity to provide prion-like properties, predicted by PLAAC^30^, span residues 13-28 and 147-214. These subsequences are comprised within a compositionally bias low-complexity region (LCR), identified by using SEG and CAST algorithms^31^, whose composition is: 18% proline, 13% alanine, 13% serine and 12% glutamine distributed within imperfect tandem repeats (P, A and Q) and homorepeats (S and A) (Fig. S1A). By performing a proline to glutamine substitution analysis using COILS algorithm^32^, an intrinsic tendency of hCPEB3’s PLD to form coiled-coils, abrogated by the Q>P substitutions, was found (Fig. S1B). hCPEB3 also contains two stretches that are computationally predicted, by AmylPred 2^33^, to form amyloids. The region with highest amyloid propensity includes residues 107-152 within the hCPEB3’s LCR (Fig. 1A), in agreement with the filament-forming segments predicted by ZipperDB algorithm (Fig. S1E)^34^. To investigate whether hCPEB3 has the ability to undergo liquid demixing, the hCPEB3 sequence was analysed with catGRANULE^35^. Whereas the whole hCPEB3 shows a low LLPS tendency, with a predicted score of 0.51 and propensity scores below the values for proteins corroborated experimentally to undergo liquid-liquid demixing (Fig. S1C)^36^, certain segments, in particular residues 254-288, show a high tendency for condensate formation (Fig. S1D).

**Figure 1.**
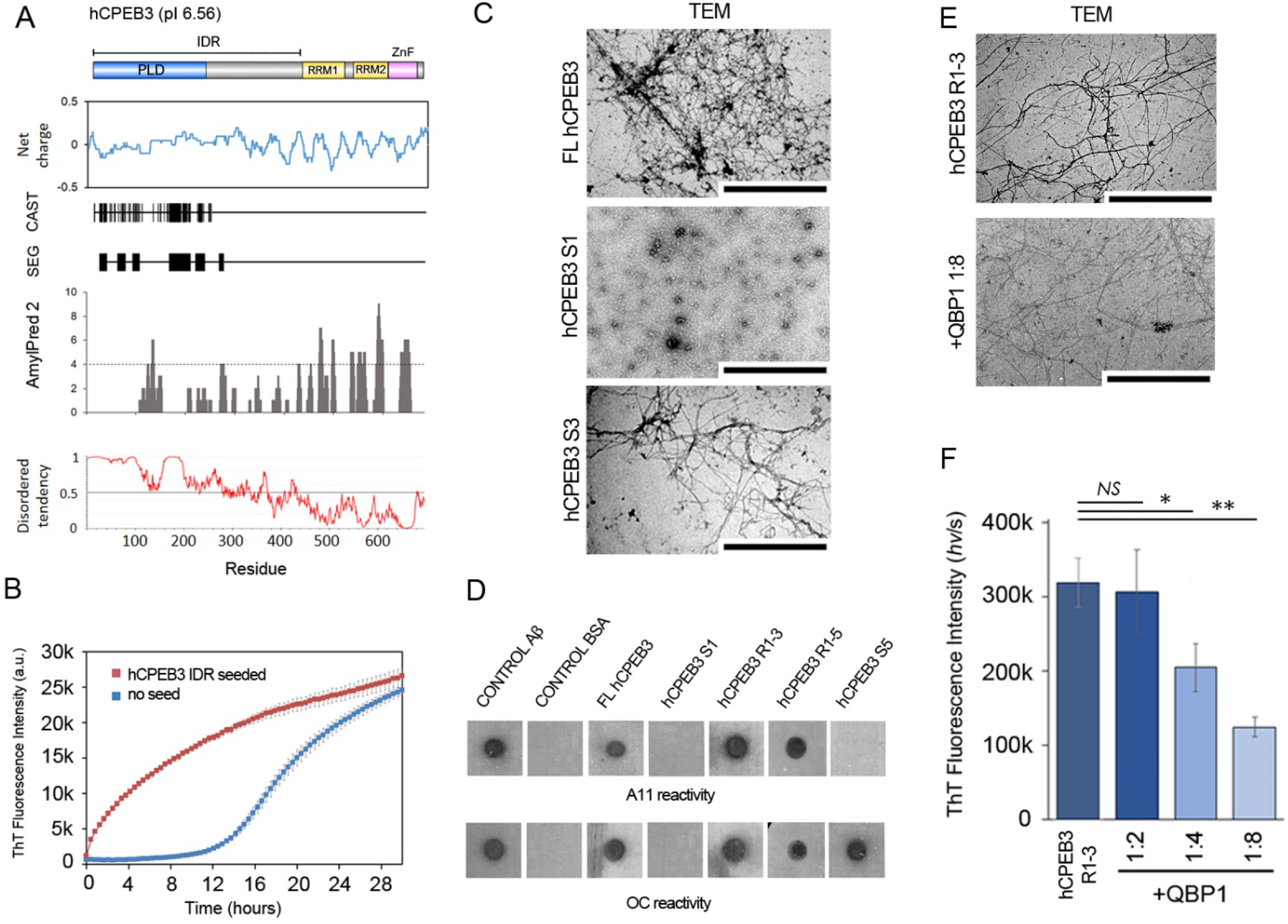
hCPEB3 IDR contains an LCR with amyloidogenic properties: **(A)** hCPEB3 representation showing the IDR and the location of the PLD (blue), RRMs (yellow) and ZnF (pink). SEG and CAST algorithms indicate that several short low complexity segments are located at the N-terminus. hCPEB3 fibrillation propensity profile as computed by the AmylPred 2 tool (dash line: minimal number of consensus programs for a residue to be considered amyloidogenic). The disorder tendency was predicted by IUPred. The position of each amino acid residue is plotted along the x-axis. **(B)** Time-course of soluble seed-free hCPEB3 IDR fibrillation in the absence (blue) and in the presence of 20% seeds (red). Error bars indicate standard error of the mean S.E.M. (n=3). **(C)** Representative electron micrograph of FL hCPEB3 amyloid filaments (scale bar: 2 *μ*m), CPEB3 S1 (1-100aa) oligomers (scale bar: 1 μm) and CPEB3 S3 (101-200aa) amyloid filaments (scale bar: 200 nm). **(D)** Immunoblot shows hCPEB3 or indicated regions and segments recognized by the A11 and OC antibodies. **(E)** Representative TEM images of hCPEB3 R1-3 (1-200aa) in absence or in presence of QBP1 in the molar ratio CPEB3:QBP1 1:8. **(F)** ThT fluorescence emission values of CPEB3 R1-3 incubated alone or with increasing concentrations of the QBP1 anti-amyloidogenic peptide. Error bars indicate S.E.M. of 3 independent experiments. *P* values were determined using Student’s *t* test: *NS*, not significant; *, *P* < 0.05; **, *P* < 0.01.

To experimentally assess the amyloidogenic behaviour of hCPEB3 *in vitro*, an aggregation-prone sequence analysis was performed by dividing the protein into four different regions and eight 100-residue overlapping segments (S1–S8), covering the whole IDR (Table S1). CPEB3’s IDR adopted an amyloid state with efficient self-templating capacity, as measured by a <u>Th</u>ioflavin-<u>T</u> (ThT) binding assay (Fig. 1B). However, the full-length (FL) hCPEB3 mainly produced amorphous aggregates with a low ThT binding capacity (Fig. S2A), accompanied by a scarce formation of mature amyloid fibers (Fig. 1C). Particularly, S1 segment (residues 1-100), which includes the Q_4_RQ_4_ polyQ tract, was found to adopt an oligomeric state (Fig. 1C). The oligomeric state adopted by S1, which was not recognized by the anti-oligomer A11 or the anti-amyloid filaments OC conformational antibodies (Fig. 1D), did not show ThT binding capacity (Fig. S2A). These data indicate that the Q_4_RQ_4_ tract is not sufficient to drive amyloid formation. The S3 segment (residues 101-200) showed an amyloid-prone behaviour, producing unbranched amyloid filaments (Fig. 1C), as predicted by both AmylPred 2 and Zipper DB algorithms. The assembled filaments exhibited the ThT binding kinetics common to amyloid-forming proteins (Fig. 2D). Also, the S3 segment, when included within R1-3 (residues 1-200) or R1-5 (residues 1-300) regions, exhibited A11 and OC reactivity (Fig. 1D). Considering that proline residues are often excluded from *β*-sheet secondary structure and that proline substitution, at any particular position, precludes amyloid formation^37–39^, we conclude that residues 106-165 are essential for hCPEB3 amyloid formation. Segment S5 (residues 202-300), which includes a poly-Ala motif, showed a high propensity to aggregate forming immature amyloid filaments (Fig. 2A). These filaments were recognized by the OC, but not the A11 antibodies (Fig. 1C), and did not show ThT binding in the same conditions (Fig. S2A).

**Figure 2.**
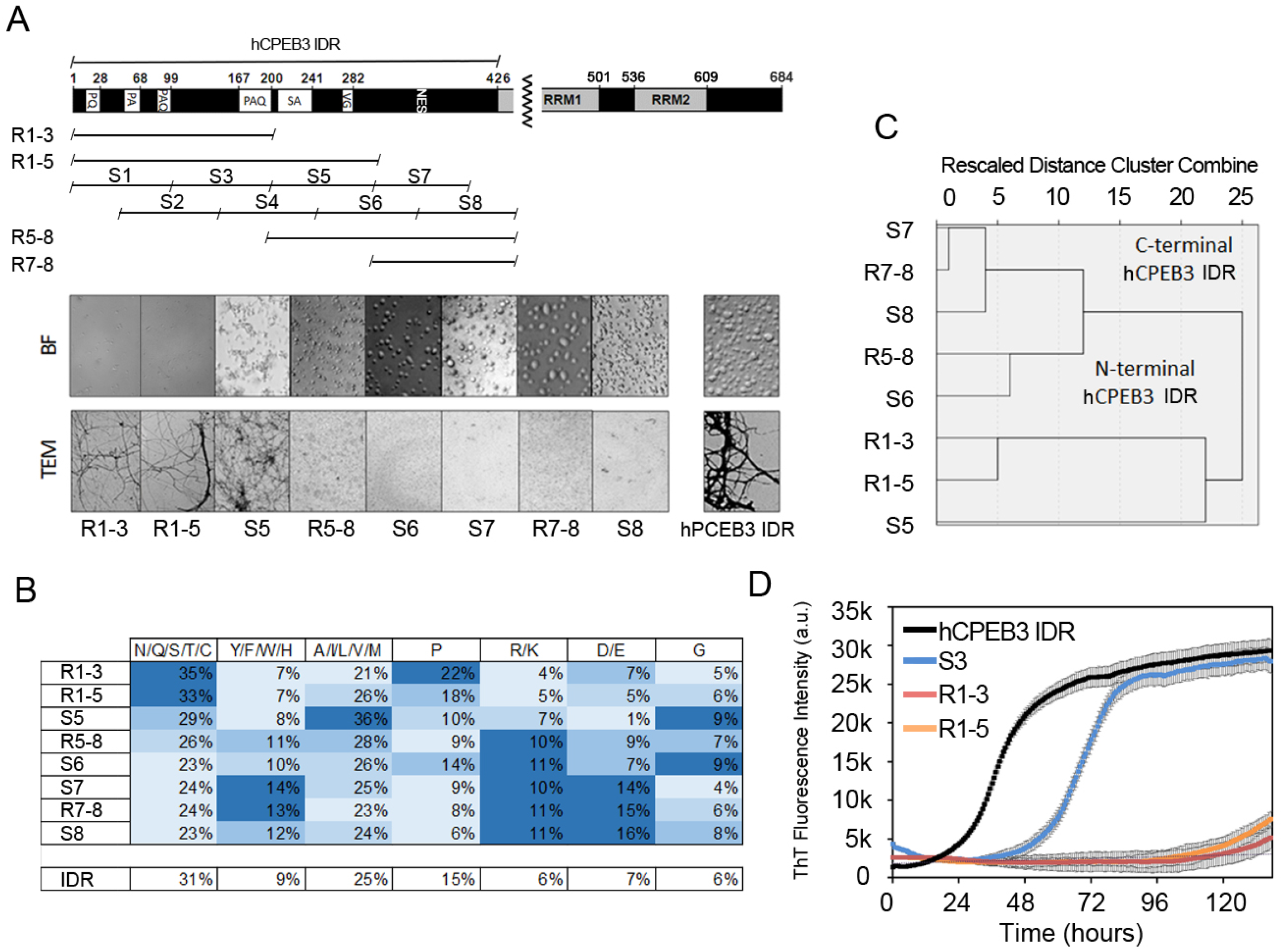
Sequential deletion analysis of hCPEB3 reveals the LLPS propensity domain: **(A)** Domain architecture of hCPEB3 and scheme of hCPEB3 recombinant segments used in this study (from S1 to S8). Representative bright field and electron micrographs show the liquid droplet formation and amyloid fibrillation of the different hCPEB3 segments assayed. **(B)** Frequency of polar, aromatic, hydrophobic, positive, negative, proline and glycine residues for each hCPEB3 sequence. **(C)** Dendrogram using average linkage between groups from hierarchical cluster analysis showing the distance or dissimilarity between clusters. **(D)** ThT fluorescence kinetics during amyloid fibrillation of hCPEB3 IDR, segment S6 and indicated regions. Data points are mean values and error bars indicate the S.E.M.

The poly<u>Q</u> binding <u>p</u>eptide <u>1</u> (QBP1) has been previously shown to block the amyloid formation of several pathological amyloid-forming proteins, such as expanded polyQ tracts^40,41^ and a TDP-43 Q/N-rich segment^42^, as well as Q/N-rich functional amyloids, such as Sup35^41^ and hCPEB3 homologs from *Aplysia* (*Ap*CPEB)^16^ and *Drosophila melanogaster* (Orb2)^13^. However, QBP1 does not affect hydrophobic amyloids like A*β*_42_^13,41^, suggesting differences in the underlying assembly mechanisms^41,43^. QBP1 reduced the amount of hCPEB3 R1-3 amyloid formation, as gauged by ThT fluorescence (Fig. 1E) and TEM (Fig. 1F), which suggests that QBP1 prevents hCPEB3 amyloid assembly likely targeting the PLD. In particular, the fact that QBP1 was raised against polyQ tracts suggests that the Q_4_RQ_4_ stretch, located at hCPEB3’s N-terminus, is essential for hCPEB3 amyloid formation.

### Residues 254-426 are necessary and sufficient for hCPEB3 liquid demixing

Next, we asked which hCPEB3’s IDR regions mediate LLPS. Previous work on RNA-binding proteins showed that phase separation requires LCRs^44–47^. Furthermore, although proline-rich LCRs, such as those of tau, polyU binding protein 1 and poly-Ala binding protein 1 were reported as not required for liquid demixing, they seem to modulate LLPS^36,48,49^. For hCPEB3 IDR, the deletion of the proline-rich R1-3 region or the poly-Ala-containing R1-5 region, both containing amyloid-forming elements, did not affect phase separation properties (Fig. 2A). All regions and segments with LLPS propensity, except for the S6 segment (residues 251-350), showed a <u>l</u>ower <u>c</u>ritical <u>s</u>olution <u>t</u>emperature point (LCST, Fig. S3A), above which condensation occurs. This observation suggests that hydrophobic interactions are the main driving forces for condensation^50^. Particularly, the hCPEB3 R5-8 region induced clustering as well as network-forming of the phase-separated droplets that ceased to grow through coalescence when compared to the other LLPS-driven elements (Fig. S3A). The hCPEB3 S6 segment showed a dual-phase behaviour corresponding to a closed loop phase diagram with an <u>u</u>pper <u>c</u>ritical <u>s</u>olution <u>t</u>emperature (UCST) higher than its LCST point (Fig. S3A). This segment, which includes residues 254-288, was identified as a liquid demixing determinant by the catGRANULE algorithm, biased towards proteins with UCST points and can explain this particular dual behaviour. Interestingly, the NES motif was not necessary for liquid demixing, given that both S6 (residues 252-350) and S8 segments (residues 352-450) undergo LLPS even though the NES motif is split in both segments. Taken together, we conclude that the hCPEB3 subsequences required for LLPS span residues 254-426 and do not display amyloidogenic properties, as the N-terminal region of the PLD does under the same conditions (Fig. S2E and S2F).

Then, the 20 amino acid residues present in proteins were grouped into seven classes (polar, hydrophobic, aromatic, positive, negative, proline and glycine) to examine the compositional distribution of each class within the segments studied here (Fig. 2B). A hierarchical cluster analysis unveiled that whereas the composition of the assayed sequences corresponding to the N-terminal LCR are divergent, those of the LLPS domain, with a composition apparently unbiased and exhibiting liquid-like properties, were redundant (P ≤ 0.05) (Fig. 2C). In this sense, as reported for other hydrophobic proteins^51^, the early droplet maturation of hCPEB IDR R5-8 correlates with a hydrophobic content increase, opposite to other regions and segments with a slower liquid-to-solid phase transition (Fig. S3A). These data suggest that residue composition, but no specific LCR sequences, can accurately predict phase transition behaviour^50^. In addition, the hCPEB3’s IDR amyloid formation kinetics monitored by ThT fluorescence, revealed that condensation, opposite to LLPS domain deletion, favours hCPEB3 amyloidogenesis (Fig. 2D). Likely, the increment in the local protein concentration within the liquid phase leads hCPEB3’s PLD to adopt the amyloid state more efficiently.

### Hydrophobic interactions are essential yet insufficient for hCPEB3 phase separation and amyloid formation

Freshly prepared, soluble recombinant hCPEB3 IDR protein in PBS pH 7.4 showed no measurable turbidity at 4 °C. However, upon raising the temperature to 37 °C, spontaneous formation of liquid droplets was observed in the absence of crowding agents like polyethylene glycol (Fig. 3A and 3B). This effect is consistent with the LCST point, where hydrophobic interactions play a key role^50^. The role of ionic strength on hCPEB3 IDR liquid droplet formation is not specific to chloride, as a similar behaviour was observed when NaF, instead of NaCl, was used in PBS (Fig. S3B). hCPEB3 IDR exhibited a temperature-dependent and reversible liquid droplet formation, as monitored by bright-field microscopy (Fig. S3C). Due to the stabilization of multivalent interactions, the liquid droplets evolved into SDS-sensitive hydrogels (Fig. S3D), as occurs for other proteins^44^. Upon further incubation, the hCPEB3 IDR hydrogels produced starbursts of ThT-positive, amyloid aggregates (Fig. S3E), similar to FUS (Fig. 3A-C)^52^. Both LLPS and amyloidogenesis processes showed a dependence with ionic strength (Fig. 3D, E). Furthermore, the poly-Ala tract included in S5, was more sensitive to the effect of the ionic strength on amyloidogenesis than the hydrophobic core included in S3. This observation positions the poly-Ala tract as the main aggregation-promoting element during hCPEB3 PLD amyloid assembly (Fig. 3E, Fig. S2C and S2D).

**Figure 3.**
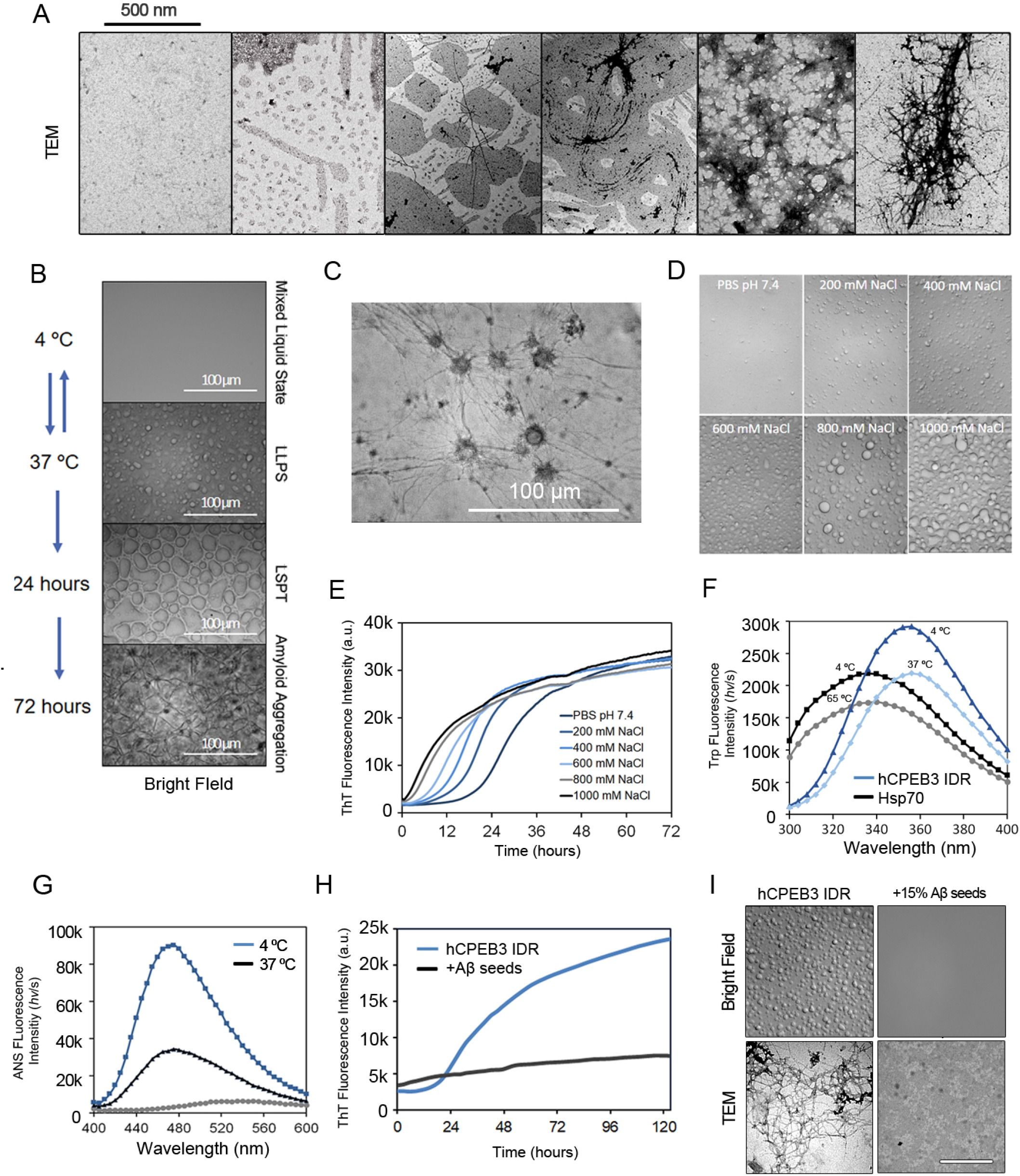
Biophysical characterization of hCPEB3 liquid-demixing: Representative **(A)** TEM and **(B)** bright field images of the morphological changes in liquid droplets of 50 *μM* hCPEB3 IDR during an “aging” experiment over 72 hr. **(C)** Morphology of hCPEB3 starburst conversion from droplets to fibrous aggregates. **(D)** Effect of the ionic strength on hCPEB3 IDR LLPS tendency visualized by bright field microscopy. Scale bar: 100 *μ*m. **(E)** ThT fluorescence kinetic curves of CPEB3 IDR showing the effect of ionic strength on amyloid formation. Each data point represents the average of three replicates. **(F)** Fluorescence emission spectra showing the effect of the temperature on the intrinsic tryptophan fluorescence of hCPEB3 IDR and Hsp70. **(G)** ANS fluorescence emission of hCPEB3 IDR in the monomeric state at 4°C and condensate at 37 °C or denatured (grey). **(H)** Effect of 15% A*β* filament seeds on hCPEB3 IDR aggregation monitored by ThT fluorescence. **(I)** Representative bright field and TEM images showing the effect of A*β* filament seeds on hCPEB3 IDR condensation and amyloidogenesis. Scale bar: 500 nm.

To gain insight into the structural properties associated to LLPS, circular dichroism (CD) spectroscopy measurements of hCPEB3’s IDR were then performed. The CD spectra of hCPEB3’s IDR recorded at 4 °C showed a minimum near 200 nm, characteristic of a statistical coil (Fig. S3F). This characteristic feature reveals a lack of abundant, stable hCPEB3’s IDR secondary structure, as was predicted *in silico* (Fig. 1A). Upon heating at 37 °C, the CD minimum signal weakened over time under high ionic strength, likely due to a decreased amount of soluble hCPEB3 IDR. hCPEB3 IDR contains five tryptophan (Trp/W) residues: W242, W252, W259 and W317, clustered within the LLPS domain, and W111 embedded within the amyloidogenic sequence. The possible presence of a stable hydrophobic core in hCPEB3 IDR was tested by measuring intrinsic Trp fluorescence, whose emission λ_max_ is < 350 nm when Trp is solvent-exposed and <340 nm when placed in a non-polar milieu. The intrinsic hCPEB3 IDR Trp fluorescence, as opposed to buried Trp residues in Hsp70 chaperone used for comparison, showed an emission λ_max_ >350 nm (Fig. 3E), which indicates the lack of a hydrophobic core in hCPEB3 IDR. The presence of exposed hydrophobic clusters was explored utilizing 8-anilinonaphthalene-1-sulfonate (ANS), which exhibits fluorescence enhancement and blue shift to 470 nm upon interaction with hydrophobic surfaces^53^. The spectral changes associated with LLPS indicated the dispersion and/or the burial of hydrophobic patches (Fig. 3G). Stretches of hydrophobic residues in the C-terminal half of A*β*_42_ have been implicated in promoting pathological aggregation in Alzheimer’s disease^54^. Thus, the effect of preformed A*β*_42_ seeds on hCPEB3 IDR liquid droplet formation was evaluated. Unexpectedly, the A*β*_42_ amyloid seeds abrogated LLPS formation and subsequent amyloid formation (Fig. 3H, I). Due to the hydrophobic nature of A*β*_42_, this inhibitory effect may be attributed to the perturbation of a hypothetical interaction between hydrophobic patches of hCPEB3 PLD with the N-terminus Q-rich tract. These results suggest that the interaction between the hydrophobic residues and the Q_4_RQ_4_ stretch is key for amyloid formation.

Hsp70/Hsp90 chaperones are known to prevent A*β*_42_ and *α*-synuclein amyloid formation *in vitro*^55^. In addition, the ability of Hsp70 to interact with exposed hydrophobic patches of substrates has been reported.^56^. Therefore, next we examined the possible effect of the Hsp70 in hCPEB3 liquid demixing and amyloidogenesis. Incubation of recombinant Hsp70 with hCPEB3’s IDR at 37 °C resulted in LLPS abrogation (Fig. 4A). To this end, an interaction with the poly-Ala tract is not required, as reflected by the results obtained with the R7-8 region, which lacks the poly-Ala tract (Fig. 4A). Itself, when included within the R1-5 region, this highly hydrophobic segment mediated PLD polymerization into higher-order filaments (Fig. S3G). Turbidity measurements at 405 nm showed that Hsp70 is able to prevent hCPEB3 IDR liquid droplet formation at equimolar concentrations (Fig. 4B) as well as the amyloid formation with sub-equimolar concentrations of Hsp70, both in the presence or absence of ATP (Fig. 4C). These results were confirmed by TEM and immuno-dot blot assays (Fig. 4D).

**Figure 4.**
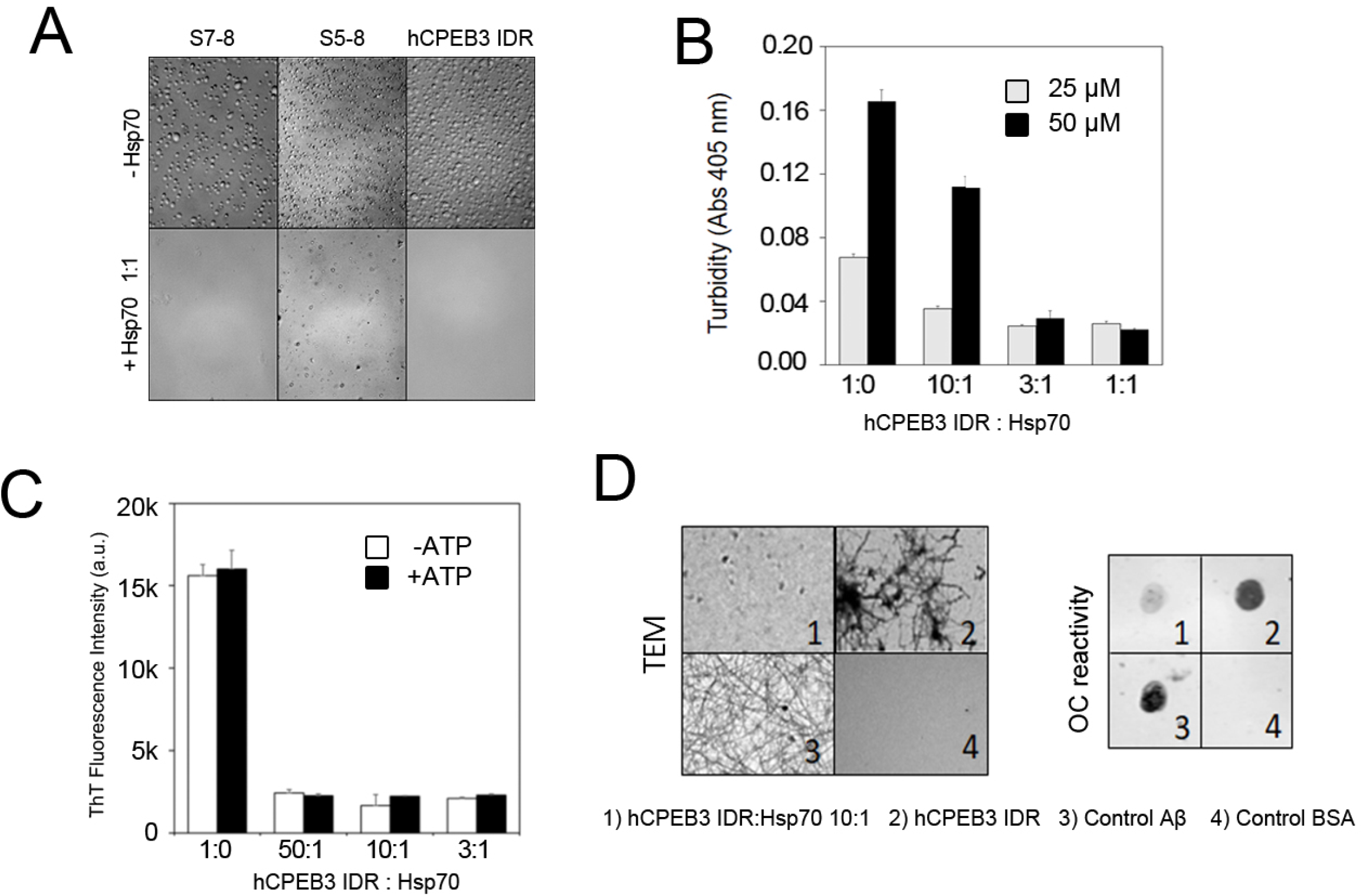
Effect of the Hsp70 chaperone on hCPEB3 liquid droplet and amyloid formation: **(A)** Representative bright field images of 50 *μ*M CPEB3 IDR liquid droplet formation and indicated regions in the absence or presence of recombinant Hsp70 chaperone. The used molar ratio was 1:1. **(B)** Liquid droplet formation of hCPEB3 IDR monitored by turbidimetry at 405 nm in the absence or in the presence of increasing concentrations of Hsp70 chaperone. Data are represented as the mean values and error bars show S.E.M. **(C)** End point values with error bars indicating S.E.M. show the effect of increasing concentrations of Hsp70 chaperone on 25 *μ*M hCPEB3 IDR amyloid filament formation followed by ThT fluorescence emission in the absence or in the presence of 2 mM ATP. **(D)** Representative electron micrographs and immuno-dot blot analysis by using the OC antibody of hCPEB3 IDR amyloid aggregation in the absence or the presence of Hsp70 at 10:1 molar ratio.

### hCPEB3, in contrast to other functional amyloids, forms long-lived toxic oligomers

Functional and pathological amyloids form toxic, metastable oligomer species^13,57^. In functional amyloids, toxic conformers are short-lived and evolve rapidly to mature, less toxic amyloid filaments within hours^13,57^. Contrary, in pathological amyloids these toxic conformers are long-lived, lasting weeks^58,59^. Similar to other amyloids, the formation of toxic oligomeric species during hCPEB3 IDR amyloidogenesis was observed by using the A11 conformational antibody (Fig. 5A). However, in contrast to other functional amyloids, such as Pmel17^60^, *Ap*CPEB^15^, Orb2^13^ and Sup35^57^, the metastable hCPEB3 IDR toxic oligomers were long-lived, similar to pathological amyloids such as A*β*_42_^61^ (Fig. 5A). Electron microscopy revealed the presence of hCPEB3 IDR annular oligomers, which eventually formed laterally associated clusters (Fig. 5B), as described for other amyloids like TDP-43^62^, *α*-synuclein^63,64^, A*β*_42_^65^, transthyretin^66^ or bacterial RepA-WH1^67^. The presence of annular oligomers was further confirmed by the *α*APF conformational antibody^68^ (Fig. 5A). We also showed that hCPEB3 IDR oligomeric species exerted cytotoxicity in neuron-like SH-SY5Y cells through necrosis (Fig. 5C and 5D)^69^. Hsp70 did not attenuate cytotoxicity once hCPEB3 IDR oligomers were formed; however, it prevented cell death when incubated with monomeric hCPEB3’s IDR (Fig. 5E). At sub-stoichiometric concentrations of Hsp70, sufficient to prevent hCPEB3 IDR amyloid formation, cytotoxicity was not abrogated until LLPS is prevented (Fig. 5E). These results suggest that toxic oligomer formation is promoted by liquid demixing and likely does not share the same pathway as the amyloid assembly.

**Figure 5.**
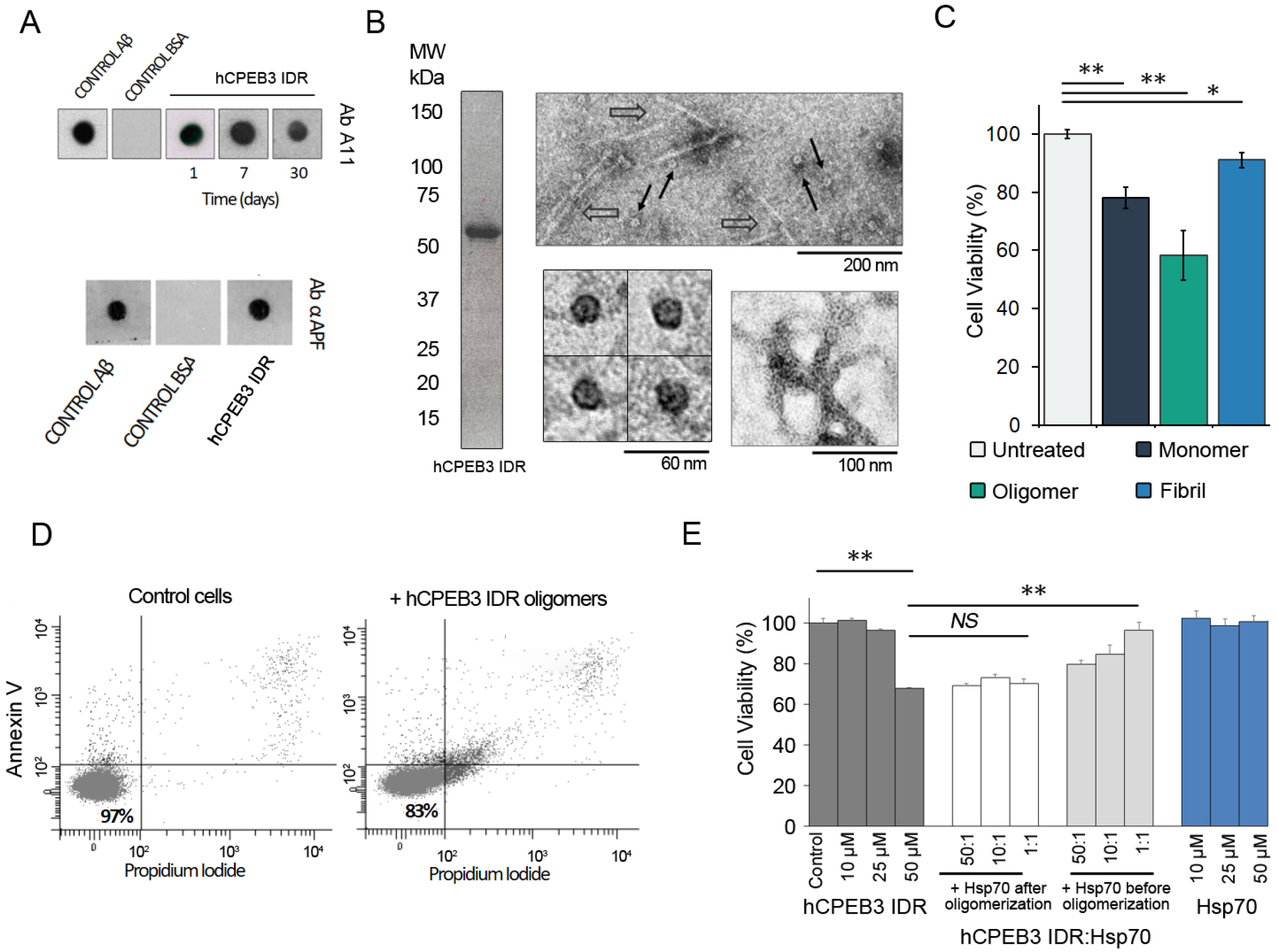
hCPEB3 shares common features with pathological amyloids: **(A)** Immuno-dot blot analysis showing the presence of stable hCPEB3 IDR toxic oligomers recognized by the A11 antibody and annular oligomers probed by the anti-APF antibody. **(B)** Coomassie blue SDS-PAGE of the freshly prepared monomeric hCPEB3 IDR and TEM images of hCPEB3 IDR annular oligomers (asterisks) and protofilaments (arrows) in the soluble fraction. Lower left panel shows the annular oligomer morphology. Lower right panel shows structural details of lateral association of annular oligomers. **(C)** Cell viability differences in SH-SY5Y cells treated with recombinant hCPEB3 IDR in the monomeric state, soluble oligomers or amyloid filaments. *P* values were determined using Student’s *t* test; *, *P* < 0.05; **, *P* < 0.01. **(D)** Flow cytometry analysis of the necrosis and apoptosis in SH-SY5Y cells exposed to hCPEB3 IDR toxic oligomers. **(E)** Effect of Hsp70 on cytotoxicity of hCPEB3 IDR oligomeric species. *P* values were determined using Student’s *t* test; *NS*, not significant.

### hCPEB3’s granules show liquid-like behaviour in neuroblastoma SH-SY5Y cells

Finally, the hCPEB3 ability to undergo phase transition in living cells was examined by overexpressing full length hCPEB3 (FL hCPEB3) and IDR in SH-SY5Y cells. FL hCPEB3, tagged with the enhanced green fluorescence protein (EGFP-FL hCPEB3), phase separated into cytoplasmic granules with irregular shapes. In contrast, EGFP-hCPEB3 IDR formed spherical granules (Fig. 6A). Live-cell imaging was used to determine whether these cytoplasmic hCPEB3 assemblies have liquid-like properties. Fusion events were frequently observed for EGFP-hCPEB3 IDR during a monitorization period of 60 s (Fig. 6B) (Movie S1A). On the other hand, neither fusion events nor net movement of the granules was detected for EGFP-FL hCPEB3 in the same timescale (Movie S1B). To better characterize the nature of hCPEB3 assemblies, FRAP experiments were performed for both the EGFP-FL hCPEB3 and the EGFP-hCPEB3 IDR. The fast FRAP recovery rates (0.25 ± 0.1 s^-1^ for EGFP-hCPEB3 IDR and 0.172 ± 0.005 s^-1^ for EGFP-FL hCPEB3) indicate that the assembled granules behave like liquids and that the EGFP-hCPEB3 IDR granule is less viscous. Both are in constant exchange with the surrounding cytoplasm (Fig. 6C). These results show that hCPEB3 phase separate into liquid-like microdroplets within SH-SY5Y cells. The observed differences between both constructions suggest that the RNA-recognition domains, and most likely their binding to mRNA, are not required for the self-assembly, but rather rigidify and stabilize the droplets.

**Figure 6.**
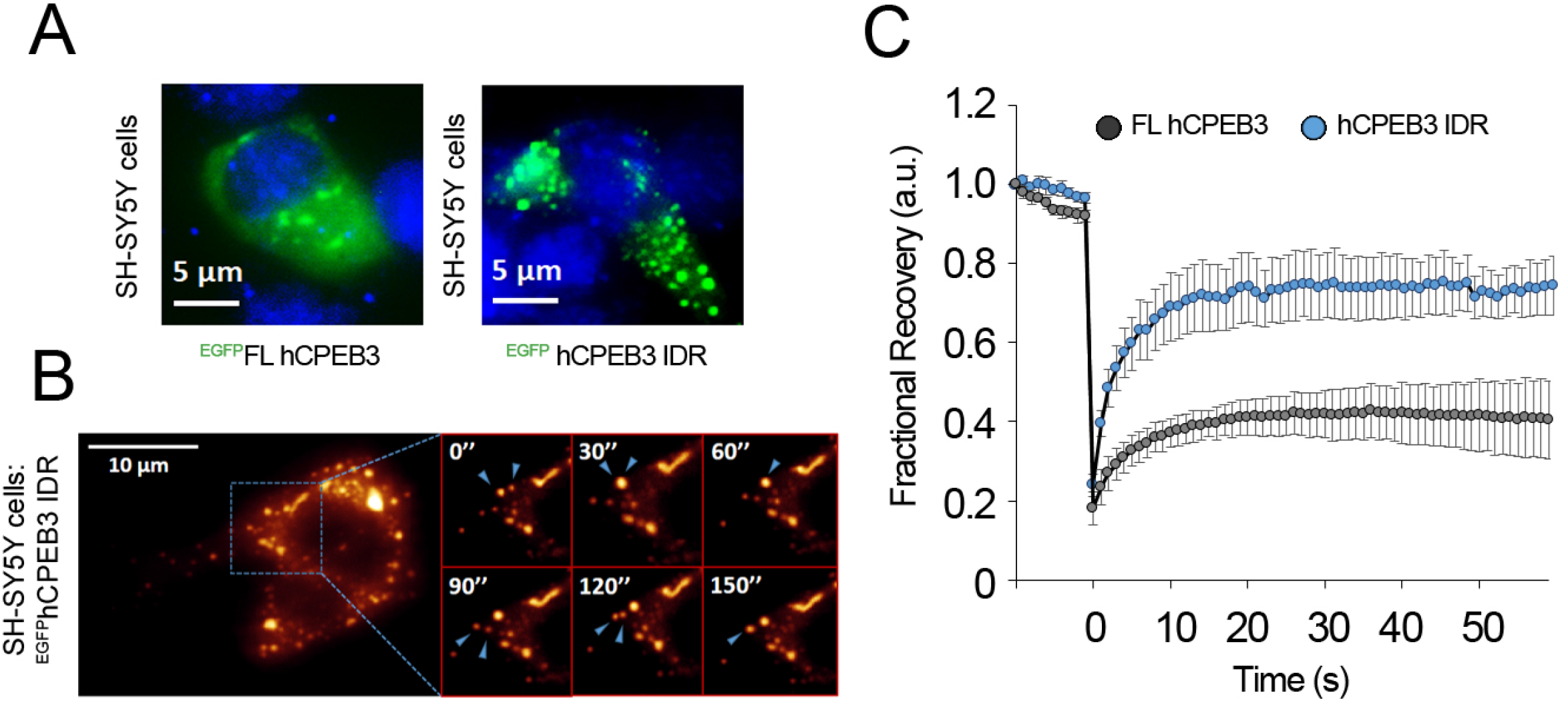
Liquid-like behaviour of hCPEB3 cellular granules in living SH-SY5Y cells: **(A)** Representative fluorescence micrographs of living SH-SY5Y cells after transient transfection overexpressing^EGFP^FL hCPEB3 or^EGFP^hCPEB3 IDR. Nuclei are shown stained with Hoechst in blue. **(B)** Representative fluorescence micrographs of fusion events of cytoplasmic^EGFP^hCPEB3 IDR droplets in living SH-SY5Y cells. A higher magnification is shown in the right panels. The two arrows point to two separated droplets before undergoing fusion, while one arrow points to the fused droplet. **(C)** FRAP experiments of^EGFP^FL hCPEB3 or^EGFP^hCPEB3 IDR. The raw data (dots) and average recovery curves (line) are shown. The error bars represent the S.E.M.

## Discussion

How long-lasting memories endure protein turnover is still unclear. However, prion-like proteins like CPEB, have the ability to adopt a self-sustaining state that would withstand the molecular turnover. Indeed, some members of the CPEB family of proteins adopt a functional amyloid state that mediates mRNA translation for long-term memory consolidation^18^. Here, by using a “divide and conquer” strategy we have identified the key sequence elements of the hCPEB3 that drive phase separation, oligomerization and amyloid aggregation. Our findings are in line with previous results from the mCPEB3, which reported that the N-terminal PLD is essential for oligomerization and amyloid formation^6^. We have also evaluated the amyloidogenic properties of the hCPEB3 *in vitro*, showing that this shares common features with pathological amyloids, which suggests that there are no intrinsic biophysical differences between them in terms of cell toxicity. Indeed, as proposed previously for FUS^52^ or TDP-43^70^, CPEB3 liquid-like condensation promotes the assembly of more stable structures by lowering the free-energy barrier of nucleation that causes toxicity. However, functional amyloids are finely tuned by evolution in order to avoid cytotoxicity^71^. Given the production of long-lasting toxic species and amyloid aggregation during hCPEB3 demixing *in vitro*, how cells avoid its cytotoxicity remains unclear^27^. Considering the well-known interplay between chaperones and IDRs^72^, we have studied the effect of Hsp70 chaperone on hCPEB3 condensation and aggregation. Based on our own results with Hsp70, and results by others with JJJ2^73^, we speculate that chaperone machinery may play a key role in regulating functional amyloidogenesis. We provide evidence showing the N-terminus is dispensable for hCPEB3 condensation and that the RNA-recognition domains are required for both hCPEB3’s cellular localization and stabilization, which supports the idea that CPEB3 can form macromolecular assemblies due to the contribution of multiple domains^24^. It has been shown that RNA-recognition domains of disease-linked RNA-binding proteins contribute to amyloid formation^74^. These observations contrast with the fact that RNA prevents liquid droplet formation in other proteins, including FUS and TDP-43^75^. Our results support the idea that phase transitions can be modulated, an observation that could be relevant as an important pharmacological target in pathological conditions. Moreover, we show that A*β*_42_ amyloid blocks hCPEB3 condensation, which further suggests that hCPEB3 could be trapped in pathological aggregates at the onset of Alzheimer disease. This could impede normal RNA translational regulation ending in memory deficits even before neurodegeneration may arise. The observations that aggregates of huntingtin with polyQ expansions trap Orb2^13^, the *Drosophila* CPEB homolog, and reduce translation has recently led to a similar proposal^76^.

### An arrangement of repetitive motifs governs hCPEB3 liquid demixing

hCPEB3 liquid-droplet formation differs from other previously reported RNA-binding proteins in which liquid demixing at physiological conditions does not require its LCR for phase separation^50^. Instead, hCPEB3 condensate formation is promoted by hydrophobic interactions that does not require the poly-Ala tract for liquid demixing, as Hsp70 chaperone can abrogate LLPS passively. Hsp70 itself possess two short intrinsically disordered LCRs with a high hydrophobic content, spanning residues 365-409 and 611-641^77^. Based on this, we suggest that rather than its C-terminal substrate-binding domain, these low-complexity tracts may block hCPEB3 condensation. The presence of segments with local LCST and UCST points within the hCPEB3 sequence reflects the idea that IDPs with LLPS tendency can be described as block-copolymers, as recently reported for Tau^78^. Thus, we propose that these different segments may enable hCPEB3 to interact with several distinct classes of condensates and coordinate their functions. We have shown how the balance between the hydrophobic and polar residues modulate the liquidity of the droplets providing important insights to understand the molecular grammar by which hCPEB3 is able to globally phase separate as well as form amyloid locally in terms of sequence. On the other hand, phosphorylation of serine residues has been reported to control biomolecular condensate formation and dissociation in FUS, a protein whose domain architecture and function share similarities with hCPEB3^79^. Since the putative phosphorylation sites of hCPEB3 IDR fall in the region identified in this work as the LLPS driver, we propose that phosphoserine residues may regulate the phase transition of hCPEB3. The components of biomolecular condensates have been divided into two qualitative classes based on their tendency to condensate: scaffolds and clients^19^. We have demonstrated that hCPEB3 spontaneously condensates in physiological conditions by itself, arising as a potential scaffold of specific RNA granules that participate in memory-related mRNA transport and repression, in contrast with the role as an apparent client recently proposed for mCPEB3^24^.

### Poly-Ala and polyQ stretches mediate hCPEB3 amyloid formation

Some IDPs contain A and Q homorepeats that tend to form high-order multimers and aggregates associated with 12 and 9 protein misfolding diseases, respectively^80^. A- and Q-rich stretches can also co-occur in the same protein, and occasionally are observed along with QA repeats of variable length, *e.g*. polyglutamine-repeat protein PQN-41, Ataxins 2 and 7, and Runt-related transcription factor 2. Above a threshold, Poly-Ala and polyQ repeats associate forming coiled-coil structures that can trigger protein aggregation and exert toxicity^81^. Although toxicity-mediated by Q-rich motifs may be important not only in pathology but also in physiological functions^82^, a variety of elements that strictly control and regulate functional LLPS and amyloid formation operate in higher organisms to avoid cytotoxicity. One example is presented in TDP-43 liquid demixing, which is mediated by a conserved A-rich hydrophobic region spanning residues 319-341^83^. Remarkably, poly-Ala tracts are conserved in mammalian CPEB3s, but they are absent in lower organisms. These observations suggest that, in the case of hCPEB3, the poly-Ala element provides an additional level of regulation for the formation of functional amyloid in mammals. A similar polar to hydrophobic content variation occurs between Ataxins 2 and 7, and their homologs in *Drosophila*^84^. Interestingly, Ser-rich regions flaking the poly-Ala tract display high prion behaviour as predicted by the PLAAC algorithm like in the case of other non-mammalian CPEB3s, which is abrogated when the poly-Ala stretch gets inserted. This observation is in agreement with previous work showing that serine is overrepresented in PLDs of human proteins linked to disease but shows a low α-helix forming propensity^47,85^. Considering that some IDPs with liquid droplet formation trends can be described as block-copolymers^86^, we propose that the polar LCRs along the hCPEB3 sequence (*i.e*., 17-QQQQRQQQQ-25, 106-SFGSTWSTGTTN-117, 136-FQQNF-140 and 236-SSASSSWNTHQS-247) would be able to adopt a cross-*β* spine or a kinked-*β* structure key for amyloid formation by cation-π and π-π interactions, as previously found for other IDPs with PLDs^87^. Finally, it must be noted that proline is a residue overrepresented within CPEB3. Considering the recent evidences support that the polyQ and proline-rich regions can drive liquid-like assembly mediated by weak hydrophobic interactions, or even aggregation with glutamine expansions as in the case of Synapsin-1^88^ and the Huntingtin Exon1^89^, and that stretches of consecutive proline residues in hCPEB3 form a PPII helix^29^, we propose that, in the case of the mammalian CPEB3s, whose amyloid state is required for biological function, proline residues would act as breakers at the Q-rich domain by limiting the amyloidogenesis of regions with prone-aggregation trends that could be harmful for neurons when being part of liquid-like assemblies.

### A model for hCPEB3’s biological function based on its structural properties

Proteins with PLDs in yeast and other organisms have recently been linked to non-toxic processes, such as the formation of stress granules and an increasing list of many other membraneless biomolecular condensates^90^. As early as 1899, cell biologist E.B. Wilson was the first to consider the cytoplasm densely packed with liquid “coacervates”. It is now well established that many cellular compartments may form in the absence of lipid membranes through LLPS processes driven by proteins, nucleic acids, and other biomolecules^91^. In this study, we have found that hCPEB3 encodes the ability to spontaneously condensate and form amyloid filaments *in vitro* in a similar fashion than other prion-like proteins. However, a fundamental aspect that makes hCPEB3 a unique and attractive system is that different “specialized” sequences are responsible for the LLPS and the amyloid-formation (Fig. 7A). Here, by combining results from our own work and previous studies, we advance a model for the biological function of hCPEB3. In the basal state, SUMOylation would block the preamutre amyloid aggregation of the PLD of hCPEB3 in agreement with the recent report of its inhibitory role in mCPEB3^92^. In this scenario, hCPEB3 target mRNAs are translationally repressed within P-body-like granules, whose formation is driven by the newly identified LLPS domain located at the C-terminus of the IDR (Fig. 7B). After neuronal stimulation, the activation of specific neuronal kinases phosphorylates hCPEB3 LLPS domain and the P-body-like granules are dissolved exposing the mRNAs to the polysomes. Once hCPEB3 is deSUMOylated, the amyloid-forming domain would adopt an active amyloid state mediated by the hydrophobic stretch and polyQ interactions, which could trigger the translation of target mRNAs and the formation of new memories within the hippocampus (Fig. 7B)^18,93^.

**Figure 7.**
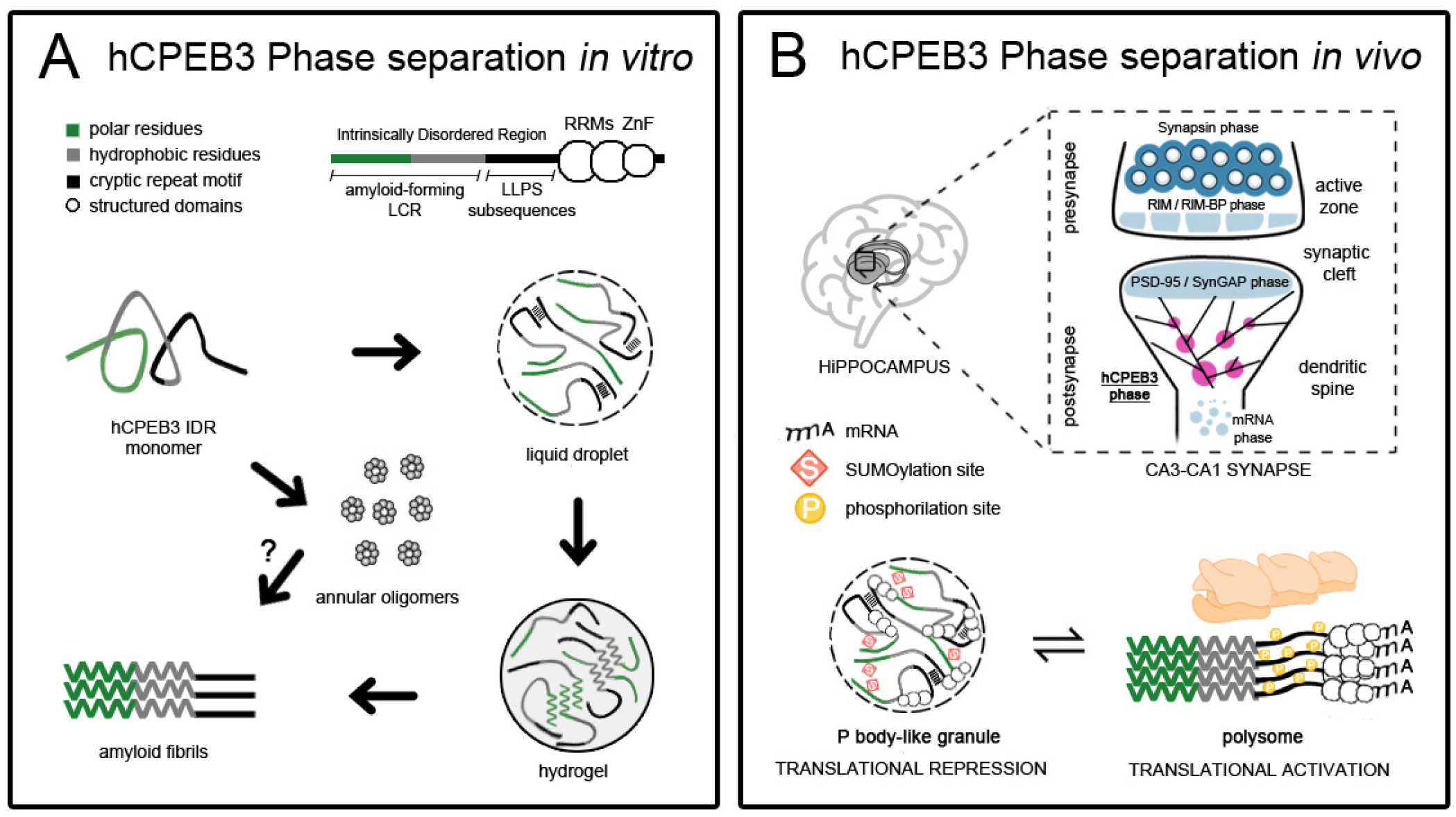
Structural basis of memory consolidation mediated by hCPEB3: **(A)** *In vitro*, recombinant hCPEB3 IDR remains stable in the monomeric state at 4 °C. When temperature increases, liquid-liquid phase separation occurs driven mainly by hydrophobic interactions among residues within the hCPEB3. Over time, liquid-to-solid phase transition turn them into hydrogels that starburst converts into to amyloid filaments formed cooperatively between hydrophobic and polyQ interactions within the amyloid-forming low-complexity region that can be blocked by A*β*_42_ and QBP1, respectively. Liquid demixing is accompanied by the formation of toxic, metastable annular oligomers. **(B)** *In vivo*, long-term memory consolidation occurs at the synaptic level within the hippocampus by local protein synthesis. In the basal state, repressed CPEB3 is SUMOylated within P body-like granules, membraneless organelle subcompartments that act as translational repressors. Upon neuronal stimulation, a decrease of SUMOylation within the amyloid-forming domain and an increase of phosphorylation within the LLPS domain dissolve P body-like granules and convert hCPEB3 into an amyloid active form that promotes the translation of target mRNAs that mediate memory storage.

## Methods

### Bioinformatics tools

Except for the PLD domain, all the other domains were predicted by SMART (http://smart.embl-heidelberg.de/) or the NCBI conserved domain (https://www.ncbi.nlm.nih.gov/Structure/cdd/wrpsb.cgi). The PLD was identified using PLAAC (http://plaac.wi.mit.edu/). The minimal contiguous PLD length for the hidden Markov model was set to 60 and the background frequencies from *Saccharomyces cerevisiae* were set to 100%. The domain structures of the proteins were generated using Illustrator for Biological Sequences. The disorder tendency was predicted by IUPred algorithm (http://iupred.enzim.hu/). To plot the sliding net charge, the sliding window was set to 20 residues by EMBOSS Bioinformatics Tool (http://www.bioinformatics.nl/cgi-bin/emboss/charge). The web platform LCR-eXXXplorer was used to detect data for low-complexity regions in protein sequences by SEG and CAST algorithms (http://repeat.biol.ucy.ac.cy/lcr-exxxplorer). AmylPred 2 combines ten different prediction tools. A peptide is considered amyloidogenic if the consensus of at least four out of ten methods is reached (http://aias.biol.uoa.gr/AMYLPRED2/). The catGRANULE algorithm predicts the tendency to assemble into foci using RNA binding and structural disordered propensities as well as amino acid patterns and the polypeptide chain (http://service.tartaglialab.com/grant_submission/catGRANULE). The COILS algorithm compares a sequence to a database of known parallel two-stranded coiled-coils and derives a similarity score (https://embnet.vital-it.ch/software/COILS_form.html). Fibrillation propensities were computed using a structure-based ZipperDB algorithm that uses the crystal structure of the fibril-forming peptide NNQQNY from the Sup35 prion protein (https://services.mbi.ucla.edu/zipperdb/). Steric zippers are predicted to form when the Rosetta energy of a hexapeptide is below the empirically determined highly negative fibrillation propensity threshold of −23 kcal/mol.

### Cloning, protein expression and protein purification

Each segment of hCPEB3 was cloned into a pET-28a vector after PCR amplifying the hCPEB3 FL gene from the pLL3.7 plasmid kindly provided by Dr. Yi-Shuian Huang^94^. The DNA amplified fragments (Table S1) were digested with *Xho*I and *Nhe*I restriction enzymes (NEB). Expression of the resulting genes led to fusion proteins containing a His6 tag and a TEV Niα protease cleavage site. All overlapping segments were expressed in the *E. coli* BL21 Star (DE3) strain using the T7 expression system (Novagene). Briefly, cells were grown in 1 L of LB medium at 37 °C shaking at 280 rpm until reaching optical densities at 595 nm (OD_595_) of ~ 0.6-0.7. Then, the cells were added of 1 mM isopropyl-β-D-thiogalactopyranoside (IPTG) for 4 hours at 37 °C to induce protein expression. Harvested cells at 4 °C, 6000 rpm (F14-6×250y rotor) for 10 min were lysed under denaturing conditions in buffer 50 mM Na_2_HPO_4_, 500 mM NaCl, 50 mM imidazole, 6 M guanidinium chloride (GdmCl) pH 7.4 with gentle agitation for 30 min and sonicated at room temperature. The lysates were centrifuged at 18000 rpm (F21-8×50y rotor) for 45 min and supernatants were purified by Ni^2+^-affinity chromatography (FPLC system ÄKTA Purifier, GE Healthcare) eluting the protein in 50 mM NaPO_4_, 500 mM NaCl, 500 mM imidazole, 3 M CdmCl, pH 7.4 while monitoring UV absorbance at 280 nm (OD_280_). Purified recombinant protein segments were diluted to ~5 *μ*M in PBS, 1 M GdmCl pH 7.4, dialysed against PBS pH 7.4 at 4 °C and then concentrated in Amicon Ultra Filters (Millipore) by centrifugation at 4000 x *g* (4 °C). Samples were stored at −80 °C until use.

The plasmid pPROEX containing the human Hsp70 gene was kindly provided by Prof. José María Valpuesta (CNB-CSIC) and transformed in the *E.coli* strain C41 (DE3). Protein expression of Hsp70 and FL hCPEB3 was induced in medium with 1 mM IPTG at 30 °C overnight. Cell lysis was carried out in non-denaturing conditions with Buffer A: 50 mM Na_2_HPO_4_, 500 mM NaCl, 50 mM imidazole, 2 mM MgCl_2_, 1% Triton X-100, 0.5% Tween 20, 1 mg/ml lysozyme, 5 *μ*g/ml DNAse I, 5 *μ*g/ml RNAse A pH 7.4 and protease inhibitor cocktail set III EDTA-free (Calbiochem) added of 1 mM PMSF. Buffer B: 50 mM NaH_2_PO_4_/Na_2_HPO_4_, 500 mM NaCl, 500 mM imidazole pH 7.4 was used as the elution buffer for His-tag affinity purification.

### TEM

A 10 *μ*L aliquot of each sample was added to formvar stabilized with carbon copper grids (Ted Pella, Inc.) for 2 minutes. The grids were negatively stained with 10 *μ*L of aqueous uranyl acetate 1% (w/v) solution for 2 minutes and then washed for two times with 10 *μ*L of water. Stain excess was removed with filter paper and the grids were air-dried. Analysis was performed at 80 kV of excitation voltage using a JEOL JEM 1200 EX II transmission electron microscope and images were recorded digitally by the SIS Megaview III CCD camera system. TEM images of 50 *μ*M hCPEB3 FL amyloid fibrils and 50 *μ*M hCPEB3 S1 oligomers were taken after 72 hours of incubation. Images of 250 *μ*M hCPEB3 S3 amyloid fibrils after 144 hours. Images of 50 *μ*M hCPEB3 R1-3 and 10 *μ*M hCPEB3 R1-5 amyloid fibrils after 216 hours. Images of 100 μM hCPEB3 S5 amyloid fibrils after 240 hours and of 50 *μ*M hCPEB3 IDR amyloid fibrils after 72 hours. The buffer used was PBS pH 7.4 and the incubation temperature was 37 °C without stiring.

### Fluorescence assays

One mM stock solution of ThT (Sigma) was prepared in PBS buffer pH 7.4. For inhibition experiments with QBP1, a 2 mM stock solution of QBP1 (Sigma) in DMSO was prepared and stored at −80 °C. The aggregation reaction was performed with 25 *μ*M hCPEB3 R1-3 at 25 °C in PBS pH 7.4 and in the presence of 2% DMSO. ThT fluorescence emission values were recorded after 4 weeks of fibril formation. 100-*μ*L samples contained 5 *μ*L of ThT (50 *μ*M). Spectra were recorded at 25 °C on a Jobin-Yvon Fuoromax-4 instrument using 3 nm excitation and emission slit widths. The excitation wavelength was 440 nm and emission was recorded over 430-500 nm at a scan speed of 1.0 nm·s^-1^. Reported values are the means of the intensities of the ThT fluorescence emission at 480 nm. The experiments were carried out in triplicate. Fluorescence intensity of ThT-monitored kinetic curves were recorded by a FLUOstar OPTIMA Microplate Reader in PBS pH 7.4 at 25 °C or in the presence of NaCl using the plate mode with excitation and emission filters of 440 nm and 490 nm, respectively, with 10 *μ*M ThT. For seeding experiments, amyloid fibrils were sonicated for 15 minutes. The concentration of hCPEB3 IDR was 50 *μ*M for kinetic experiments unless indicated otherwise. Intrinsic tryptophan fluorescence spectra of 25 *μ*M hCPEB3 IDR and Hsp70 were recorded varying the temperature from 4 °C to 37 °C or 65 °C, respectively, between 300 and 400 nm at 280 nm excitation in PBS pH 7.4. Two mM of ANS (Sigma) stock solution was prepared in PBS pH 7.4. ANS fluorescence emission spectra of 25 *μ*M hCPEB3 in either monomeric state at 4 °C or condensate at 37 °C PBS 1 M NaCl pH 7.4 were recorded by using a Jobin-Yvon Fuoromax-4 fluorometer. Denaturing conditions were PBS 3 M GdmCl pH 7.4. The final ANS concentration in samples was 40 *μ*M. The spectra were recorded as the accumulation of three consecutive scans and the conditions of the acquisition were: excitation wavelength of 350 nm, emission range from 400 to 600 nm. A Peltier module was used to control the temperature.

### Dot blot assay

Two *μ*L aliquots of each sample were spotted onto a nitrocellulose membrane. A*β* (1-42) and BSA were used as positive and negative controls, respectively. After blocking for 45 minutes at room temperature with 10% non-fat milk (Blotting-Grade Blocker, Bio-Rad) in TBS containing 0.01% Tween 20 (TBS-T), the membrane was washed three times for 5 minutes with TBS-T and incubated for one hour at room temperature with the polyclonal specific anti-oligomer A11 antibody (ThermoScientific) diluted 1:2000, the αAPF antibody diluted 1:1000 or the fiber-specific OC antibody (Millipore) diluted 1:1000 in 5% non-fat milk in TBS-T. The membrane was washed three times for 5 minutes with TBS-T before being incubated for 1 hour with IRDye 680LT Goat anti-rabbit (LI-COR) and then revealed in an Odyssey^®^ CLx (LI-COR) apparatus. The concentrations of hCPEB3 and indicated regions and segments were as follows: [FL hCPEB3] = 50 *μ*M; [hCPEB3 IDR] = 25 *μ*M; [hCPEB3 S1] = 0.3 mM; [hCPEB3 R1-3] = 0.5 mM; [hCPEB3 R1-5] = 0.2 mM; [hCPEB3 S5] = 0.1 mM. 20 *μ*M A*β* fibrils and 60 *μ*M BSA were used as positive and negative controls respectively.

### Circular Dichroism

The far-UV CD spectra of the hCPEB3 IDR protein at a concentration of 5-10 *μ*M were collected in 5 mM Na_2_HPO_4_, 5 mM KH_2_PO_4_ pH 7.4 or with 1 M NaF with a JASCO-J810 spectropolarimeter (JASCO Inc.) equipped with a Peltier temperature control unit and using quartz cuvettes of 1 mm pathlength. Four scans were accumulated and averaged and the buffer contribution was subtracted from the experimental data. Spectra were recorded from 260 to 190 nm at the scan speed of 20 nm/min. The instrument bandwidth was 1.25 nm and the data pitch was 0.5 mm.

### Cell viability and cell death assays

The CellTiter-Glo^®^ Luminescent Cell Viability Assay (Promega) was performed in opaque-walled 96-well plates. 2 x 10^4^ neuroblastoma SH-SY5Y cells per well were allowed to attach in DMEM/F12 1:1 GlutaMAX medium supplemented with 10% FBS and 1% penicillin/streptomycin for 24 hours prior to the addition of hCPEB3 IDR samples. To obtain amyloid hCPEB3 soluble oligomers, monomeric hCPEB3 IDR was incubated in PBS pH 7.4 at 37 °C. After 24 hours, the supernatant containing soluble amyloid oligomers and the pellet fraction were separated by centrifugation at 14.000 x *g* for a short spin and collected. For fibrils, the insoluble fraction was collected after 72 hr of incubation and sonicated for 15 min. Cells were treated with 50 *μM* hCPEB3 IDR in the monomeric state, soluble oligomer fraction or amyloid fibrils. For experiments with Hsp70, hCPEB3 soluble oligomers were produced at the indicated concentrations and used at 50 *μ*M when co-incubated with the chaperone. Cells were incubated for 24 hours and then the plate was tempered to room temperature for approximately 30 minutes. One-hundred *μ*L of CellTiter-Glo^®^ Reagent was added to the same volume of medium-containing cells. Content was mixed for 2 minutes on an orbital shaker to induce cell lysis and was incubated at room temperature for 10 minutes to stabilize the signal. Luminiscence was recorded using a FLUOstar OPTIMA microplate reader.The mean of readings of triplicate wells was taken as one value. The value for the control cultures was considered as 100% viability. For Annexin V-FITC/propidium iodide assay, 8 x 10^4^ neuroblastoma SH-SY5Y cells seeded in 24-well plates were treated with 50 *μ*M hCPEB3 IDR soluble oligomers. The cells were then removed after 24 hr by trypsinization, rinsed with PBS and resuspended in binding buffer containing Annexin V and propidium iodide for 15 minutes in the dark. The samples were analysed on a BD FACSAria Cell Sorter and data were recorded using FlowJo Software.

### Constructs and SH-SY5Y cell transfection

Primers used for generating pEGFP-FL hCPEB3 1-698 (Forward primer: CCGCTCGAGGCATGCAGGATGATTTACTGATG. Reverse primer: CGGGATCCTCAGCTCCAGCGGAACGGGACGTG) and pGEFP-CPEB3 IDR 1-426 (Forward primer: CCGCTCGAGGCATGCAGGATGATTTACTGATG. Reverse primer: CGGGATCCTCAAAACCTGCGAAAGCTGGC). For FL hCPEB3, the product of PCR amplification was subcloned in a pCR2.1 plasmid (TA cloning kit, Invitrogen). The DNA insert and the hCPEB3 IDR PCR product were digested with *Xho*I and *Bam*HI restriction enzymes (NEB) and cloned into a pEGFP-C1 expression vector. Direct sequencing of the constructs confirmed successful cloning. The day before transfection, cells were seeded into 35 mm glass bottom dishes (Ibidi GmbH, Germany) at a density of 2 x 10^5^ cells. Approximately, 60-80% confluence was reached on the day of transfection. Lipofectamine 3000 (Invitrogen, CA, USA) was mixed with 2.5 *μ*g of plasmids for EGFP-FL CEPB3 (1-698aa), GFP-CPEB3 IDR (1-426aa) or empty pEGFP-C1 as control at 1:1 molar ratio with serum-free DMEM/F12 medium and DNA-lipid complexes were incubated for 15 minutes at room temperature before adding to cells uniformly. Transfected cells were incubated for 36 hours prior to analysis. Cells were visualized by using a fluorescence microscope Leica DMI 6000.Excitation filter: BP 470/40; emission filter BP 525/50; dichromatic mirror: 500.

### FRAP methods and analysis

The FRAP experiments were performed using a Leica TCS SP5 inverted confocal microscope. Time-lapse images of the samples were collected by using the argon laser at 488 nm for 1 min at 1 frame *per* second using an HCX PL APO CS 40.0X oil objective with a scan speed of 1400 Hz. For FRAP analysis, fluorescence intensities from two regions of interest (ROI) of time-lapse images were computed. ROI-1 was the photobleached droplet while ROI-2 was defined as background, and its signal was subtracted from ROI-1. The photobleaching corrected fluorescence intensity *vs*. time graphs were expressed in fractional form normalized by the pre-photobleach intensity. The relative intensity of 8 granules *per* condition from different cells of 3 different transient transfections was used to calculate the mean and the standard deviation. Images were analysed with Leica LAS AF Lite software.

## Acknowledgements

We thank Dr. Yi-Shuian Huang for providing us the pLL3.7 plasmid, Prof. José María Valpuesta for the pPROEX plasmid, Dr. Laura Barrios for assistance with statistical analysis and Prof. Rakez Kayed for the αAPF annular anti-protofibril antibody. We are also grateful for excellent technical assistance on transmission electron microscopy to Dr. Ricardo Martínez and Martin I. Maher, Dr. Silvia Fernández, Dr. Emilio Tejera and Andrea Collado of the cellular and molecular biology facility, M^a^ Carmen Hernández and M^a^ Belén García of the microscopy and scientific image facility and José Luis Martínez of the flow cytometry facility. This work was supported by grants SAF2013-49179-C2-1-R and SAF2016-76678-C2-1-R to MC-V and SAF2016-76678-C2-2-R to DVL from the Spanish Ministry of Economy and Competiveness.

## References

1. Kandel, E. R. The molecular biology of memory: CAMP, PKA, CRE, CREB-1, CREB-2, and CPEB. Molecular Brain 5, (2012).

2. Mendez, R. & Richter, J. D. Translational control by CPEB: A means to the end. Nature Reviews Molecular Cell Biology 2, 521–529 (2001).

3. Richter, J. D. CPEB: a life in translation. Trends Biochem. Sci. 32, 279–85 (2007).

4. Huang, Y. S., Kan, M. C., Lin, C. L. & Richter, J. D. CPEB3 and CPEB4 in neurons: Analysis of RNA-binding specificity and translational control of AMPA receptor GluR2 mRNA. EMBO J. 25, 4865–4876 (2006).

5. Chao, H. W. et al. Deletion of CPEB3 enhances hippocampus-dependent memory via increasing expressions of PSD95 and NMDA receptors. J. Neurosci. 33, 17008–17022 (2013).

6. Stephan, J. S. et al. The CPEB3 Protein Is a Functional Prion that Interacts with the Actin Cytoskeleton. Cell Rep. 11, 1772–85 (2015).

7. Fioriti, L. et al. The Persistence of Hippocampal-Based Memory Requires Protein Synthesis Mediated by the Prion-like Protein CPEB3. Neuron 86, 1433–48 (2015).

8. Si, K. & Kandel, E. R. The Role of Functional Prion-Like Proteins in the Persistence of Memory. Cold Spring Harb. Perspect. Biol. 8, a021774 (2016).

9. Sudhakaran, I. P. & Ramaswami, M. Long-term memory consolidation: The role of RNA-binding proteins with prion-like domains. RNA Biol. 14, 568–586 (2017).

10. Si, K., Lindquist, S. & Kandel, E. R. A neuronal isoform of the aplysia CPEB has prion-like properties. Cell 115, 879–91 (2003).

11. Shorter, J. & Lindquist, S. Prions as adaptive conduits of memory and inheritance. Nat. Rev. Genet. 6, 435–50 (2005).

12. Majumdar, A. et al. Critical role of amyloid-like oligomers of Drosophila Orb2 in the persistence of memory. Cell 148, 515–29 (2012).

13. Hervás, R. et al. Molecular Basis of Orb2 Amyloidogenesis and Blockade of Memory Consolidation. PLoS Biol. 14, (2016).

14. Fiumara, F., Fioriti, L., Kandel, E. R. & Hendrickson, W. A. Essential Role of Coiled Coils for Aggregation and Activity of Q/N-Rich Prions and PolyQ Proteins. Cell 143, 1121–1135 (2010).

15. Si, K., Choi, Y. B., White-Grindley, E., Majumdar, A. & Kandel, E. R. Aplysia CPEB Can Form Prion-like Multimers in Sensory Neurons that Contribute to Long-Term Facilitation. Cell 140, 421–435 (2010).

16. Hervás, R. et al. Divergent CPEB prion-like domains reveal different assembly mechanisms for a generic amyloid-like fold. bioRxiv 2020.05.19.103804 (2020). doi:10.1101/2020.05.19.103804

17. Cervantes, S. A. et al. Identification and Structural Characterization of the N-terminal Amyloid Core of Orb2 isoform A. Sci. Rep. 6, (2016).

18. Hervás, Rubén; J. Rau, Michael; Park, Younshim; Zhang, Wenjuan; G. Murzin, Alexey; A.J. Fitzpatrick, James; H.W. Scheres; Si, K. Cryo-EM structure of a neuronal functional amyloid implicated in memory persistence in Drosophila. Science (80-.). 376, 1230–4 (2020).

19. Banani, S. F., Lee, H. O., Hyman, A. A. & Rosen, M. K. Biomolecular condensates: organizers of cellular biochemistry. Nat. Rev. Mol. Cell Biol. 18, 285–298 (2017).

20. Woodruff, J. B., Hyman, A. A. & Boke, E. Organization and Function of Non-dynamic Biomolecular Condensates. Trends Biochem. Sci. 43, 81–94 (2018).

21. Guo, L. et al. Nuclear-Import Receptors Reverse Aberrant Phase Transitions of RNA-Binding Proteins with Prion-like Domains. Cell 173, 677–692.e20 (2018).

22. Hofweber, M. & Dormann, D. Friend or foe-Post-translational modifications as regulators of phase separation and RNP granule dynamics. Journal of Biological Chemistry 294, 7137–7150 (2019).

23. Khan, M. R. et al. Amyloidogenic Oligomerization Transforms Drosophila Orb2 from a Translation Repressor to an Activator. Cell 163, 1468–1483 (2015).

24. Ford, L., Ling, E., Kandel, E. R. & Fioriti, L. CPEB3 inhibits translation of mRNA targets by localizing them to P bodies. 116, 18078–18087 (2019).

25. Kaczmarczyk, L. et al. New phosphospecific antibody reveals isoform-specific phosphorylation of CPEB3 protein. PLoS One 11, (2016).

26. Peng, S.-C., Lai, Y.-T., Huang, H.-Y., Huang, H.-D. & Huang, Y.-S. A novel role of CPEB3 in regulating EGFR gene transcription via association with Stat5b in neurons. doi:10.1093/nar/gkq634

27. Chuang, E., Hori, A. M., Hesketh, C. D. & Shorter, J. Amyloid assembly and disassembly. J. Cell Sci. 131, (2018).

28. Dosztányi, Z., Csizmok, V., Tompa, P. & Simon, I. IUPred. Bioinformatics 21, 3433–3434 (2005).

29. Mingo, D. R. de, Pantoja-Uceda, D., Hervás, R., Vázquez, M. C. & Laurents, D. V. Preferred Conformations in the Intrinsically Disordered Region of Human CPEB3 Explain its Role in Memory Consolidation. bioRxiv 2020.05.12.091587 (2020). doi:10.1101/2020.05.12.091587

30. Lancaster, A. K., Nutter-Upham, A., Lindquist, S. & King, O. D. PLAAC: a web and command-line application to identify proteins with prion-like amino acid composition. Bioinformatics 30, 2501–2 (2014).

31. Kirmitzoglou, I. & Promponas, V. J. LCR-eXXXplorer: A WEB platform to search, visualize and share data for low complexity regions in protein sequences. Bioinformatics 31, 2208–2210 (2015).

32. Lupas, A., Van Dyke, M. & Stock, J. Predicting coiled coils from protein sequences. Science (80-.). 252, 1162–1164 (1991).

33. Tsolis, A. C., Papandreou, N. C., Iconomidou, V. A. & Hamodrakas, S. J. A Consensus Method for the Prediction of ‘Aggregation-Prone’ Peptides in Globular Proteins. PLoS One 8, (2013).

34. Thompson, M. J. et al. The 3D profile method for identifying fibril-forming segments of proteins. Proc. Natl. Acad. Sci. U. S. A. 103, 4074–8 (2006).

35. Mitchell, S. F., Jain, S., She, M. & Parker, R. Global analysis of yeast mRNPs. Nat. Struct. Mol. Biol. 20, 127–133 (2013).

36. Ambadipudi, S., Biernat, J., Riedel, D., Mandelkow, E. & Zweckstetter, M. Liquid-liquid phase separation of the microtubule-binding repeats of the Alzheimer-related protein Tau. Nat. Commun. 8, (2017).

37. Moriarty, D. F. & Raleigh, D. P. Effects of sequential proline substitutions on amyloid formation by human amylin20-29. Biochemistry 38, 1811–1818 (1999).

38. Kraus, A. Proline and lysine residues provide modulatory switches in amyloid formation: Insights from prion protein. Prion 10, 57–62 (2016).

39. Thakur, A. K., Yang, W. & Wetzel, R. Inhibition of polyglutamine aggregate cytotoxicity by a structure-based elongation inhibitor. FASEB J. 18, 923–925 (2004).

40. Nagai, Y. et al. A toxic monomeric conformer of the polyglutamine protein. Nat. Struct. Mol. Biol. 14, 332–40 (2007).

41. Hervás, R. et al. Common features at the start of the neurodegeneration cascade. PLoS Biol. 10, (2012).

42. Mompeán, M., Ramírez de Mingo, D. & Hervás, R. Molecular mechanism of the inhibition of TDP-43 amyloidogenesis by QBP1. Arch. Biochem. Biophys. (2019).

43. Nagai, Y. et al. Inhibition of polyglutamine protein aggregation and cell death by novel peptides identified by phage display screening. J. Biol. Chem. 275, 10437–42 (2000).

44. Han, T. W. W. et al. Cell-free formation of RNA granules: low complexity sequence domains form dynamic fibers within hydrogels. Cell 149, 768–779 (2012).

45. Lin, Y., Protter, D. S. W., Rosen, M. K. & Correspondence, R. P. Formation and Maturation of Phase-Separated Liquid Droplets by RNA-Binding Proteins. Mol. Cell 60, 208–219 (2015).

46. Molliex, A. et al. Phase separation by low complexity domains promotes stress granule assembly and drives pathological fibrillization. 163, (2015).

47. Aguzzi, A. & Altmeyer, M. Phase Separation: Linking Cellular Compartmentalization to Disease. Trends in Cell Biology 26, 547–558 (2016).

48. Riback, J. A. et al. Stress-Triggered Phase Separation Is an Adaptive, Evolutionarily Tuned Response. Cell 168, 1028–1040.e19 (2017).

49. Franzmann, T. M. & Alberti, S. Prion-like low-complexity sequences: Key regulators of protein solubility and phase behavior. (2018). doi:10.1074/jbc.TM118.001190

50. Martin, E. W. & Mittag, T. Relationship of Sequence and Phase Separation in Protein Low-Complexity Regions. Biochemistry 57, 2478–2487 (2018).

51. Muiznieks, L. D., Sharpe, S., Pomès, R. & Keeley, F. W. Role of Liquid–Liquid Phase Separation in Assembly of Elastin and Other Extracellular Matrix Proteins. Journal of Molecular Biology 430, 4741–4753 (2018).

52. Patel, A. et al. A Liquid-to-Solid Phase Transition of the ALS Protein FUS Accelerated by Disease Mutation Article A Liquid-to-Solid Phase Transition of the ALS Protein FUS Accelerated by Disease Mutation. Cell 162, (2015).

53. Rosen, C. G. & Weber, G. Dimer Formation from 1-Anilino-8-naphthalenesulfonate Catalyzed by Bovine Serum Albumin. A New Fluorescent Molecule with Exceptional Binding Properties. Biochemistry 8, 3915–3920 (1969).

54. Kim, W. & Hecht, M. H. Generic hydrophobic residues are sufficient to promote aggregation of the Alzheimer’s Abeta42 peptide. Proc. Natl. Acad. Sci. U. S. A. 103, 15824–9 (2006).

55. Lackie, R. E. et al. The Hsp70/Hsp90 chaperone machinery in neurodegenerative diseases. Frontiers in Neuroscience 11, (2017).

56. Clerico, E. M., Tilitsky, J. M., Meng, W. & Gierasch, L. M. How Hsp70 molecular machines interact with their substrates to mediate diverse physiological functions. Journal of Molecular Biology 427, 1575–1588 (2015).

57. Krishnan, R. et al. Conserved features of intermediates in amyloid assembly determine their benign or toxic states. Proc. Natl. Acad. Sci. U. S. A. 109, 11172–11177 (2012).

58. Bucciantini, M. et al. Inherent toxicity of aggregates implies a common mechanism for protein misfolding diseases. Nature 416, 507–11 (2002).

59. Bucciantini, M. et al. Prefibrillar amyloid protein aggregates share common features of cytotoxicity. J. Biol. Chem. 279, 31374–82 (2004).

60. Fowler, D. M. et al. Functional amyloid formation within mammalian tissue. PLoS Biol. 4, e6 (2006).

61. Bartolini, M. et al. Kinetic characterization of amyloid-beta 1-42 aggregation with a multimethodological approach. Anal. Biochem. 414, 215–225 (2011).

62. Fang, Y. S. et al. Full-length TDP-43 forms toxic amyloid oligomers that are present in frontotemporal lobar dementia-TDP patients. Nat. Commun. 5, (2014).

63. Lorenzen, N. et al. The role of stable α-synuclein oligomers in the molecular events underlying amyloid formation. J. Am. Chem. Soc. 136, 3859–68 (2014).

64. Chen, S. W. et al. Structural characterization of toxic oligomers that are kinetically trapped during α-synuclein fibril formation. Proc. Natl. Acad. Sci. U. S. A. 112, E1994–E2003 (2015).

65. Lasagna-Reeves, C. A. et al. Tau oligomers impair memory and induce synaptic and mitochondrial dysfunction in wild-type mice. Mol. Neurodegener. 6, 39 (2011).

66. Pires, R. H., Karsai, Á., Saraiva, M. J., Damas, A. M. & Kellermayer, M. S. Z. Distinct Annular Oligomers Captured along the Assembly and Disassembly Pathways of Transthyretin Amyloid Protofibrils. PLoS One 7, (2012).

67. Fernandez, C., Nuñez-Ramirez, R., Jimenez, M., Rivas, G. & Giraldo, R. RepA-WH1, the agent of an amyloid proteinopathy in bacteria, builds oligomeric pores through lipid vesicles. Sci. Rep. 6, (2016).

68. Kayed, R. et al. Annular Protofibrils Are a Structurally and Functionally Distinct Type of Amyloid Oligomer *. (2008). doi:10.1074/jbc.M808591200

69. Behl, C., Davis, J. B., Klier, F. G. & Schubert, D. Amyloid beta peptide induces necrosis rather than apoptosis. Brain Res. 645, 253–64 (1994).

70. Gasset-Rosa, F. et al. Cytoplasmic TDP-43 De-mixing Independent of Stress Granules Drives Inhibition of Nuclear Import, Loss of Nuclear TDP-43, and Cell Death. Neuron 102, 339–357.e7 (2019).

71. Jackson, M. P. & Hewitt, E. W. Why are functional amyloids non-toxic in humans? Biomolecules 7, (2017).

72. Pechmann, S. & Frydman, J. Interplay between Chaperones and Protein Disorder Promotes the Evolution of Protein Networks. PLoS Comput. Biol. 10, (2014).

73. Li, L. et al. A Putative Biochemical Engram of Long-Term Memory. Curr. Biol. 26, 3143–3156 (2016).

74. Agrawal, S. et al. RNA recognition motifs of disease-linked RNA-binding proteins contribute to amyloid formation. Sci. Rep. 9, (2019).

75. Maharana, S. et al. RNA buffers the phase separation behavior of prion-like RNA binding proteins. Science 360, 918–921 (2018).

76. Joag, H. et al. A role of cellular translation regulation associated with toxic Huntingtin protein. Cell. Mol. Life Sci. (2019). doi:10.1007/s00018-019-03392-y

77. Chakafana, Zininga & Shonhai. The Link That Binds: The Linker of Hsp70 as a Helm of the Protein’s Function. Biomolecules 9, 543 (2019).

78. Boyko, S., Qil, X., Chen, T. H., Surewicz, K. & Surewicz, W. K. Liquid-liquid phase separation of tau protein: The crucial role of electrostatic interactions. J. Biol. Chem. 294, 11054–11059 (2019).

79. Monahan, Z. et al. Phosphorylation of the FUS low-complexity domain disrupts phase separation, aggregation, and toxicity. EMBO J. 36, 2951–2967 (2017).

80. Darling, A. L. & Uversky, V. N. Intrinsic disorder in proteins with pathogenic repeat expansions. Molecules 22, (2017).

81. Pelassa, I. et al. Association of polyalanine and polyglutamine coiled coils mediates expansion disease-related protein aggregation and dysfunction. Hum. Mol. Genet. 23, 3402–3420 (2014).

82. Blum, E. S., Schwendeman, A. R. & Shaham, S. polyQ disease: misfiring of a developmental cell death program? (2012). doi:10.1016/j.tcb.2012.11.003

83. Conicella, A. E., Zerze, G. H., Mittal, J. & Fawzi, N. L. ALS Mutations Disrupt Phase Separation Mediated by α-Helical Structure in the TDP-43 Low-Complexity C-Terminal Domain. Structure 24, 1537–1549 (2016).

84. Bakthavachalu, B. et al. RNP-Granule Assembly via Ataxin-2 Disordered Domains Is Required for Long-Term Memory and Neurodegeneration. Neuron 98, 754–766.e4 (2018).

85. Chakrabartty, A., Kortemme, T. & Baldwin, R. L. Helix propensities of the amino acids measured in alanine-based peptides without helix-stabilizing side-chain interactions. Protein Sci. 3, 843–852 (1994).

86. Ruff, K. M., Roberts, S., Chilkoti, A. & Pappu, R. V. Advances in Understanding Stimulus-Responsive Phase Behavior of Intrinsically Disordered Protein Polymers. Journal of Molecular Biology 430, 4619–4635 (2018).

87. Hughes, M. P. et al. Atomic structures of low-complexity protein segments reveal kinked b sheets that assemble networks. Science (80-.). 359, 698–701 (2018).

88. Milovanovic, D., Wu, Y., Bian, X. & De Camilli, P. CELL BIOLOGY A liquid phase of synapsin and lipid vesicles.

89. Peskett, T. R. et al. A Liquid to Solid Phase Transition Underlying Pathological Huntingtin Exon1 Aggregation. Mol. Cell 70, 588–601 (2018).

90. Franzmann, T. M. et al. Phase separation of a yeast prion protein promotes cellular fitness. Science 359, (2018).

91. Alberti, S., Gladfelter, A. & Mittag, T. Considerations and Challenges in Studying Liquid-Liquid Phase Separation and Biomolecular Condensates. Cell 176, 419–434 (2019).

92. Drisaldi, B. et al. SUMOylation Is an Inhibitory Constraint that Regulates the Prion-like Aggregation and Activity of CPEB3. Cell Rep. 11, 1694–702 (2015).

93. Giese, K. P. & Mizuno, K. The roles of protein kinases in learning and memory. Learning and Memory 20, 540–552 (2013).

94. Huang, W. H., Chao, H. W., Tsai, L. Y., Chung, M. H. & Huang, Y. S. Elevated activation of CaMKIIα in the CPEB3-knockout hippocampus impairs a specific form of NMDAR-dependent synaptic depotentiation. Front. Cell. Neurosci. 8, (2014)

