## Supplemental Material for "Molecular Determinants of Liquid Demixing and Amyloidogenesis in Human CPEB3"

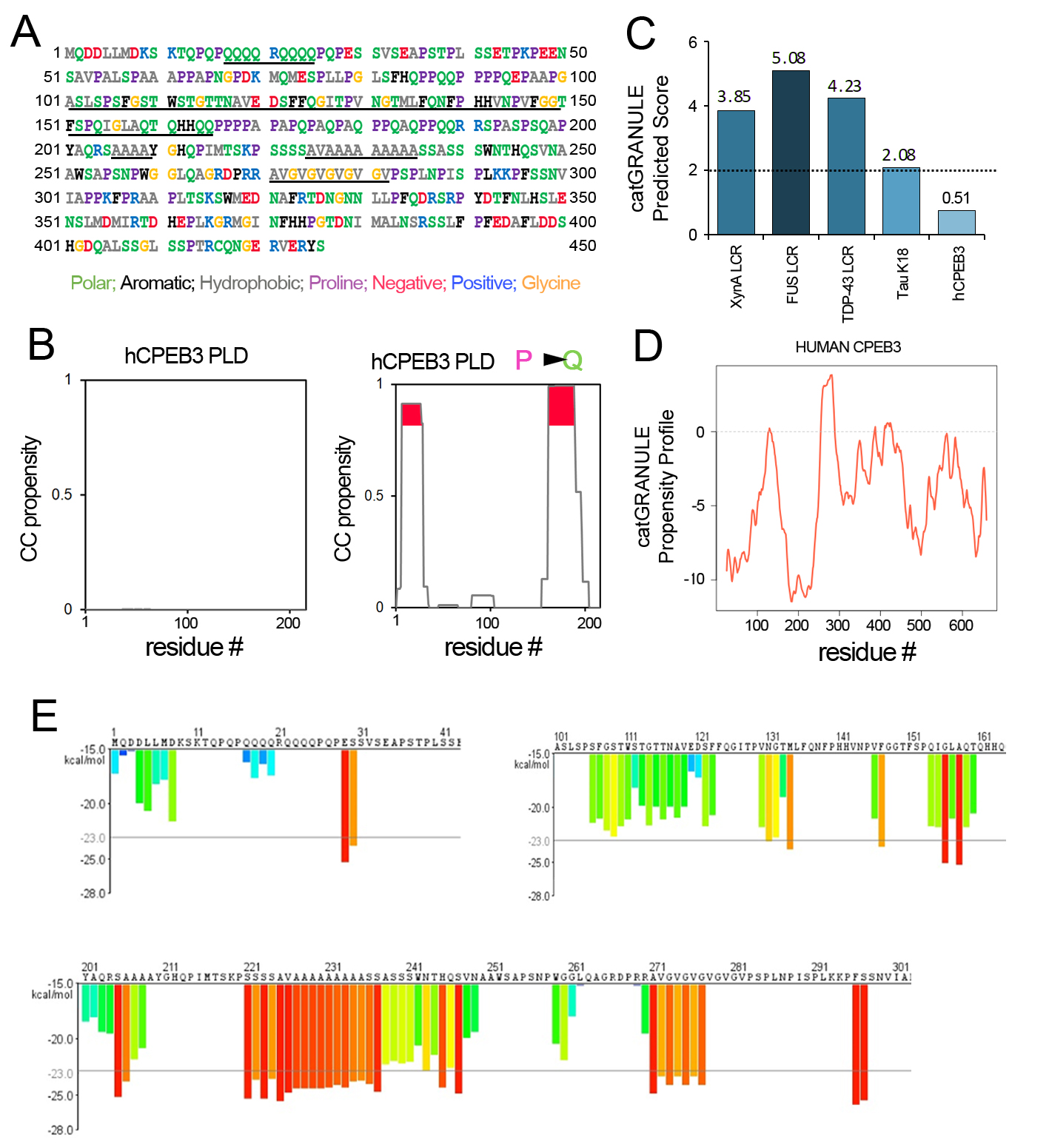

***Supplementary Figure 1. Bioinformatic Analysis of the hCPEB3 sequence:*** (A) Intrinsically disordered hCPEB3 sequence is shown with polar, aromatic, hydrophobic, proline negatively charged, positively charged and glycine residues coloured green, black, grey, fuchsia, red, blue and yellow, respectively. The key elements for hCPEB3 oligomerization and amyloid formation are shown underlined in black. (B) *Per*-residue coiled-coil (CC) propensity for hCPEB3 PLD or hCPEB3 PLD Pro to Gln substitution as calculated by COILS algorithm. Peaks with the highest propensity (0.8-1) are highlighted in red. (C) Total catGRANULE scores of different domains of experimentally corroborated liquid droplet-forming proteins and hCPEB3. (D) Residue-specific propensity score of hCPEB3 sequence for granule formation predicted by catGRANULE. (E) Fibrillation propensity of hCPEB3 to form steric zippers. The amino acid residues were coloured according to their fibrillation propensities in a rainbow scale from blue (low) to red (high).

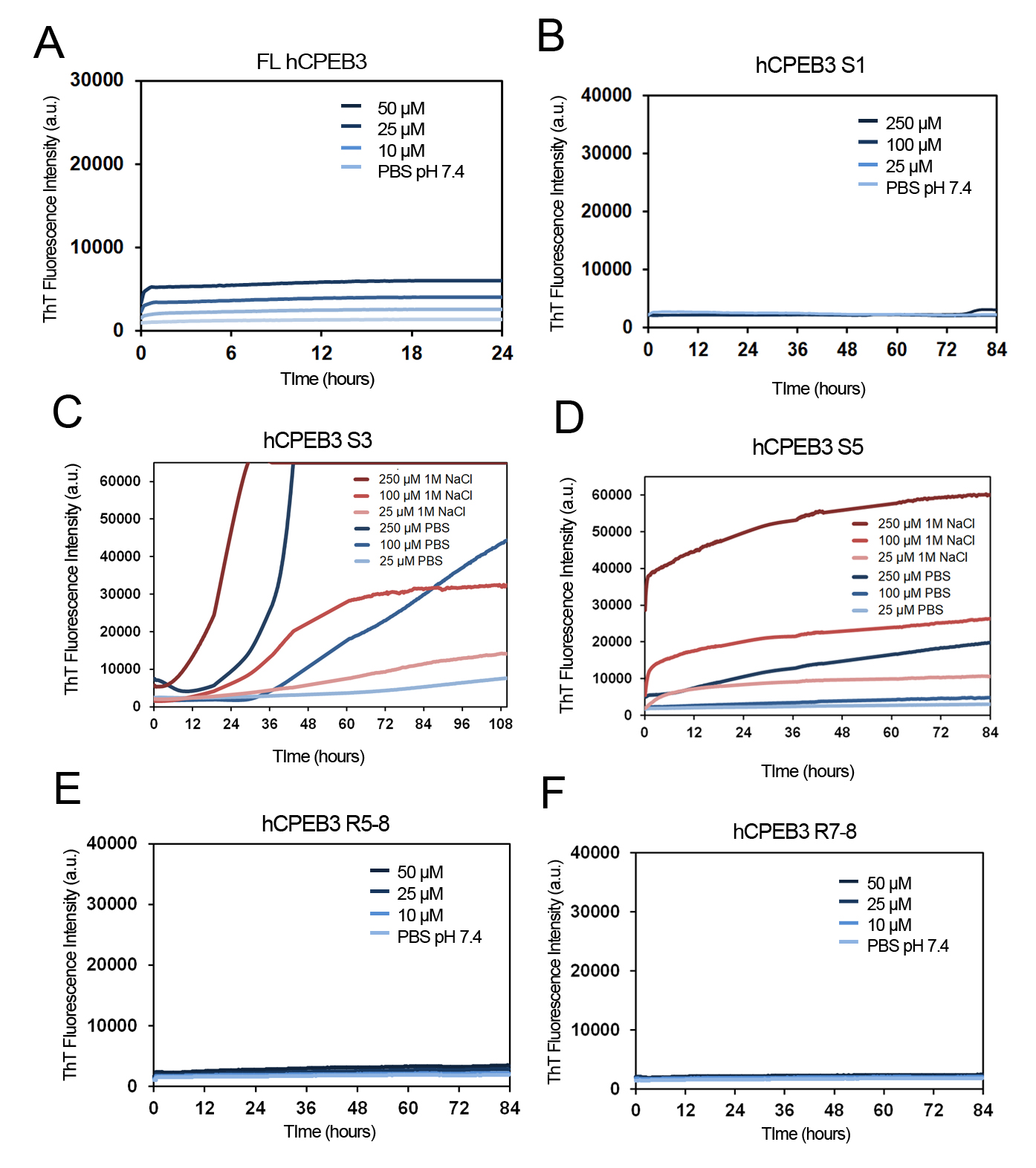

***Supplementary Figure 2. ThT analysis of hCPEB3 fibrillation kinetics:*** (A) Time-course of amyloid fibrillation followed by ThT fluorescence emission of FL hCPEB3 and (B) hCPEB3 S1 (1-100aa) at the indicated concentrations. (C) Time-course showing the effect of ionic strength on fibrillation rates for hCPEB3 S3 (101-200) segment and (D) hCPEB3 S5 (202-300). Experimental conditions are indicated in the corresponding panels. (E) Time-course of amyloid fibrillation followed by ThT fluorescence emission of hCPEB3 R5-8 (202-450aa) and (F) hCPEB3 R7-8 (302-450aa) at the indicated concentrations.

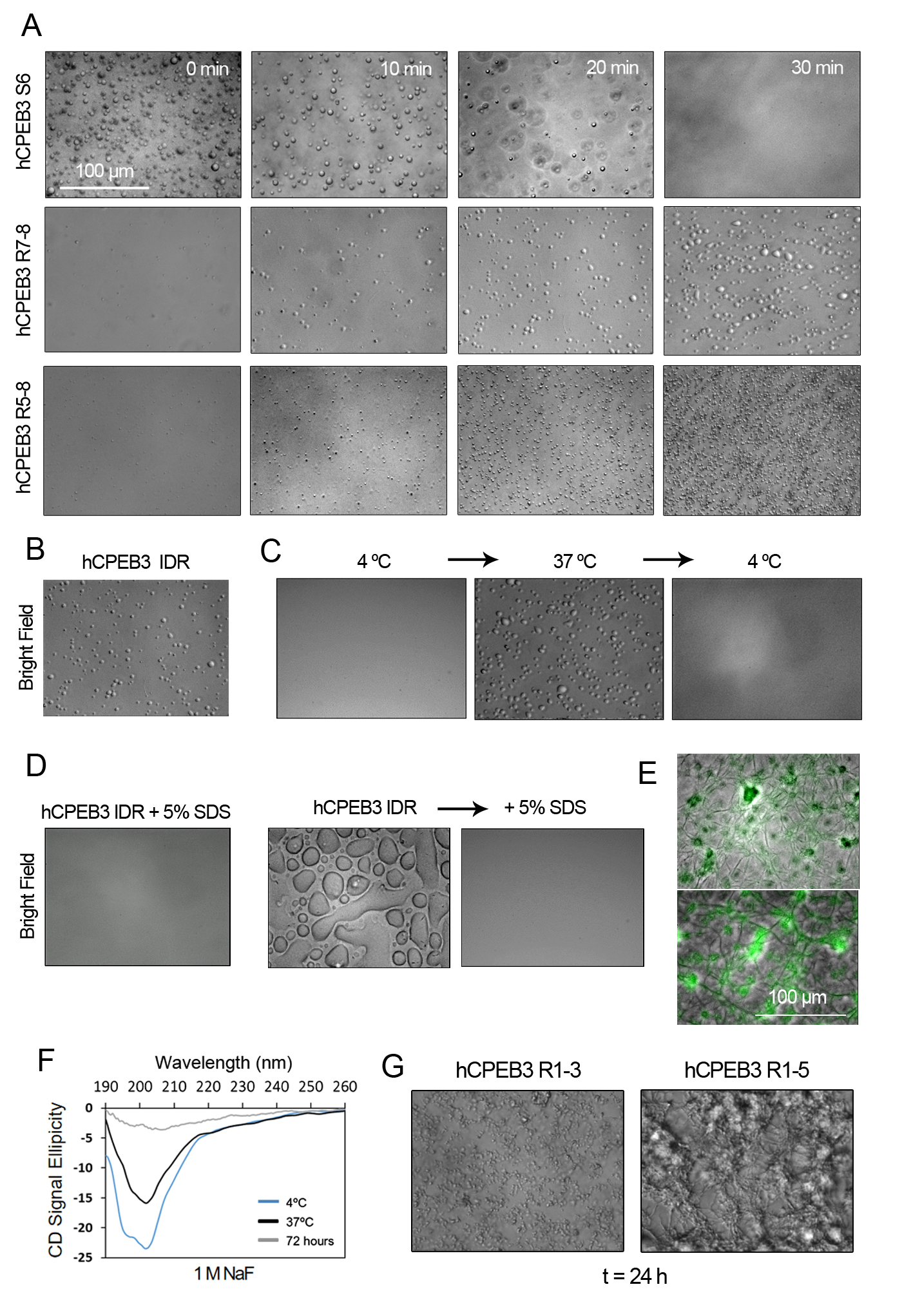

***Supplementary Figure 3. Structural properties of hCPEB3 condensation, amyloid formation and oligomerization:*** (A) Representative bright field images of hCPEB3 S6 segment, hCPEB3 R7-8 and R5-8 regions showing liquid droplet dissolution upon time. (B) Representative bright field image of hCPEB3 IDR liquid droplet formation in 10 mM NaH_2_PO_4_, 1 M NaF. (C) Effect of temperature on LLPS. hCPEB3 IDR liquid droplets were visualized by bright field microscopy. (D) The left panel shows a representative bright field image of the effect of 5% SDS on hCPEB3 IDR LLPS and the right panel shows the solubilisation effect of 5% SDS on preformed hCPEB3 IDR hydrogels. (E) hCPEB3 IDR amyloid filaments stained with ThT were visualized by fluorescence microscopy. (F) Far- UV circular dichroism spectra of 5-10 *µ*M hCPEB3 IDR in the monomeric state at 4 ºC or in the condensate state at 37 ºC, and, 72 hr after LLPS at 25 ºC. (G) Bright field representative images of hCPEB3 R1-3 (1-200aa) and R1-5 (1-300aa) oligomerization after 24 hr of incubation. [hCPEB3 S6] = 50 *µ*M; [hCPEB3 IDR] = 50 *µ*M except for CD: [hCPEB3 R1-3] = 0.5 mM; [hCPEB3 R1-5] = 0.2 mM.

***Supplementary Table 1.*** Primers and PCR conditions used to generate the plasmids containing each recombinant region or segment from hCPEB3.

| **PCR conditions** | 30 cycles | | | 30 cycles | | | 30 cycles | | | 30 cycles | | | 30 cycles | | | 30 cycles | | | 30 cycles | | | 30 cycles | | |
| --- | --- | --- | --- | --- | --- | --- | --- | --- | --- | --- | --- | --- | --- | --- | --- | --- | --- | --- | --- | --- | --- | --- | --- | --- |
|  | 10´´ | 30´´ | 15´´ | 10´´ | 30´´ | 15´´ | 10´´ | 30´´ | 15´´ | 10´´ | 30´´ | 15´´ | 10´´ | 30´´ | 15´´ | 10´´ | 30´´ | 15´´ | 10´´ | 30´´ | 15´´ | 10´´ | 30´´ | 15´´ |
|  | 98 ºC | 59 ºC | 72 ºC | 98 ºC | 60 ºC | 72 ºC | 98 ºC | 60 ºC | 72 ºC | 98 ºC | 60 ºC | 72 ºC | 98 ºC | 60 ºC | 72 ºC | 98 ºC | 60 ºC | 72 ºC | 98 ºC | 63 ºC | 72 ºC | 98 ºC | 60 ºC | 72 ºC |
|  | DEN | ANN | EXT | DEN | ANN | EXT | DEN | ANN | EXT | DEN | ANN | EXT | DEN | ANN | EXT | DEN | ANN | EXT | DEN | ANN | EXT | DEN | ANN | EXT |
| **Primer reverse sequence** | CACCTCGAGTCATCAGCCCG  GTGCCGCGGGCTCC | | | CACCTCGAGTCATCAAGTGC  CTCCGGAAGACTGGG | | | CACCTCGAGTCATCAGGGCG  CCTGGCTGGGGCTGGC | | | CACCTCGAGTCATCATGCAT  TCACGCTTTGGTGC | | | CACCTCGAGTCATCACACGT  TGCTGGAGAAGGGC | | | CACCTCGAGTCATCACTCCA  ACGAGTGCAAGTTAAAAG | | | CACCTCGAGTCATCAGCTAT  CATCCAGGAAGGC | | | CACCTCGAGTCATCAAAACC  TGCGAAAGCTGGC | | |
| **Primer forward sequence** | CTAGCTAGCGAAAACCTGTATTTTCA  GATGCAGGATGATTTACTGATGG | | | CTAGCTAGCGAAAACCTGTATTTTCA  GAGCGCAGTGCCGGCCCTCAGCC | | | CTAGCTAGCGAAAACCTGTATTTTCAGGCCTGGAGCGCACCGTCCAACC | | | CTAGCTAGCGAAAACCTGTATTTTCA  GTCCTTAATGGATATGATAAGG | | | CTAGCTAGCGAAAACCTGTATTTTCA  GTCCTTAATGGATATGATAAGG | | | CTAGCTAGCGAAAACCTGTATTTTCA  GTCCTTAATGGATATGATAAGG | | | CTAGCTAGCGAAAACCTGTATTTTCA  GGCGCCGCCCAAGTTCCCTCGC | | | CTAGCTAGCGAAAACCTGTATTTTCA  GTCCTTAATGGATATGATAAGG | | |
| **SEGMENT** | **CPEB3 S1**  **1-100aa** | | | **CPEB3 S2**  **51-150aa** | | | **CPEB3 S3**  **101-200aa** | | | **CPEB3 S4**  **152-250aa** | | | **CPEB3 S5**  **202-300aa** | | | **CPEB3 S6**  **251-350aa** | | | **CPEB3 S7**  **302-400aa** | | | **CPEB3 S8**  **352-450aa** | | |

| **PCR conditions** | 30 cycles | | | 30 cycles | | | 30 cycles | | | 30 cycles | | | 30 cycles | | |
| --- | --- | --- | --- | --- | --- | --- | --- | --- | --- | --- | --- | --- | --- | --- | --- |
|  | 10´´ | 30´´ | 1´ | 10´´ | 30´´ | 1´ | 10´´ | 30´´ | 1´ | 10´´ | 30´´ | 45´´ | 10´´ | 30´´ | 45´´ |
|  | 98 ºC | 63 ºC | 72 ºC | 98 ºC | 62 ºC | 72 ºC | 98 ºC | 62 ºC | 72 ºC | 98 ºC | 60 ºC | 72 ºC | 98 ºC | 60 ºC | 72 ºC |
|  | DEN | ANN | EXT | DEN | ANN | EXT | DEN | ANN | EXT | DEN | ANN | EXT | DEN | ANN | EXT |
| **Primer reverse sequence** | CACCTCGAGTCATCAAAACC  TGCGAAAGCTGGC | | | CACCTCGAGTCATCAGGGCG  CCTGGCTGGGGCTGGC | | | CACCTCGAGTCATCACACGT  TGCTGGAGAAGGGC | | | CACCTCGAGTCATCAGCTAT  CATCCAGGAAGGC | | | CACCTCGAGTCATCAGCTAT  CATCCAGGAAGGC | | |
| **Primer forward sequence** | CTAGCTAGCGAAAACCTGTATTTTCA  GATGCAGGATGATTTACTGATGG | | | CTAGCTAGCGAAAACCTGTATTTTCA  GAGCGCAGTGCCGGCCCTCAGCC | | | CTAGCTAGCGAAAACCTGTATTTTCAGGCCTGGAGCGCACCGTCCAACC | | | CTAGCTAGCGAAAACCTGTATTTTCA  GTCCTTAATGGATATGATAAGG | | | CTAGCTAGCGAAAACCTGTATTTTCA  GGCGCCGCCCAAGTTCCCTCGC | | |
| **SEGMENT** | **CPEB3 IDR**  **1-426aa** | | | **CPEB3 R1-3**  **1-200aa** | | | **CPEB3 R1-5**  **1-300aa** | | | **CPEB3 R5-8**  **202-450aa** | | | **CPEB3 R7-8**  **302-450aa** | | |

***Supplementary Movies.*** hCPEB3 forms dynamic compartments in SH-SY5Y cells. (A) A time-lapse movie of an EGFP-CPEB3 IDD-overexpressing living SH-SY5Y cells. (B) A time-lapse movie of an EGFP-FL CPEB3-overexpressing living SH-SY5Y cells.

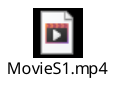
